# Fanconi Anemia Pathway Deficiency Drives Copy Number Variation in Squamous Cell Carcinomas

**DOI:** 10.1101/2021.08.14.456365

**Authors:** Andrew L.H. Webster, Mathijs A. Sanders, Krupa Patel, Ralf Dietrich, Raymond J. Noonan, Francis P. Lach, Ryan R. White, Audrey Goldfarb, Kevin Hadi, Matthew M. Edwards, Frank X. Donovan, Moonjung Jung, Sunandini Sridhar, Olivier Fedrigo, Huasong Tian, Joel Rosiene, Thomas Heineman, Jennifer A. Kennedy, Lorenzo Bean, Ozgur Rosti, Rebecca Tryon, Ashlyn-Maree Gonzalez, Allana Rosenberg, Ji-Dung Luo, Thomas Carrol, Eunike Velleuer, Jeff C. Rastatter, Susanne I. Wells, Jordi Surrallés, Grover Bagby, Margaret L. MacMillan, John E. Wagner, Maria Cancio, Farid Boulad, Theresa Scognamiglio, Roger Vaughan, Amnon Koren, Marcin Imielinski, Settara Chandrasekharappa, Arleen D. Auerbach, Bhuvanesh Singh, David I. Kutler, Peter J. Campbell, Agata Smogorzewska

## Abstract

Fanconi anemia (FA), a model syndrome of genome instability, is caused by a deficiency in DNA interstrand crosslink (ICL) repair resulting in chromosome breakage^1–3^. The FA repair pathway comprises at least 22 FANC proteins including BRCA1 and BRCA2^4–6^, and protects against carcinogenic endogenous and exogenous aldehydes^7–10^. Individuals with FA are hundreds to thousands-fold more likely to develop head and neck (HNSCC), esophageal and anogenital squamous cell carcinomas (SCCs) with a median onset age of 31 years^11^. The aggressive nature of these tumors and poor patient tolerance of platinum and radiation-based therapy have been associated with short survival in FA^11–16^. Molecular studies of SCCs from individuals with FA (FA SCCs) have been limited, and it is unclear how they relate to sporadic HNSCCs primarily driven by tobacco and alcohol exposure or human papillomavirus (HPV) infection^17^. Here, by sequencing FA SCCs, we demonstrate that the primary genomic signature of FA-deficiency is the presence of a high number of structural variants (SVs). SVs are enriched for small deletions, unbalanced translocations, and fold-back inversions that arise in the context of TP53 loss. The SV breakpoints preferentially localize to early replicating regions, common fragile sites, tandem repeats, and SINE elements. SVs are often connected forming complex rearrangements. Resultant genomic instability underlies elevated copy number alteration (CNA) rates of key HNSCC-associated genes, including *PIK3CA, MYC, CSMD1, PTPRD, YAP1, MXD4, and EGFR.* In contrast to sporadic HNSCC, we find no evidence of HPV infection in FA HNSCC, although positive cases were identified in gynecologic tumors. A murine allograft model of FA pathway-deficient SCC was enriched in SVs, exhibited dramatic tumor growth advantage, more rapid epithelial-to-mesenchymal transition (EMT), and enhanced autonomous inflammatory signaling when compared to an FA pathway-proficient model. In light of the protective role of the FA pathway against SV formation uncovered here, and recent findings of FA pathway insufficiency in the setting of increased formaldehyde load resulting in hematopoietic stem cell failure and carcinogenesis^18–20^, we propose that high copy-number instability in sporadic HNSCC may result from functional overload of the FA pathway by endogenous and exogenous DNA crosslinking agents. Our work lays the foundation for improved FA patient treatment and demonstrates that FA SCC is a powerful model to study tumorigenesis resulting from DNA crosslinking damage.

---

To profile the mechanism of tumor formation in the setting of DNA interstrand crosslink (ICL) repair deficiency, derive biomarkers for early detection, and identify potential therapeutic opportunities for individuals with FA, we sequenced 55 independent FA SCCs and three adenocarcinomas from 50 individuals. Clinical data, which was available for 41 individuals from this cohort, revealed that they developed SCCs at a young median age of 31 years and have a median cancer-specific survival of only 17 months, much shorter than patients with sporadic HPV positive or negative HNSCCs^21^ **(Fig1A and Extended data Fig1A).** Characteristics of the FA individuals and sequenced tumors are described in **Extended data Fig 1B-D** and **Supplemental Table 1**.

**Fig. 1.**
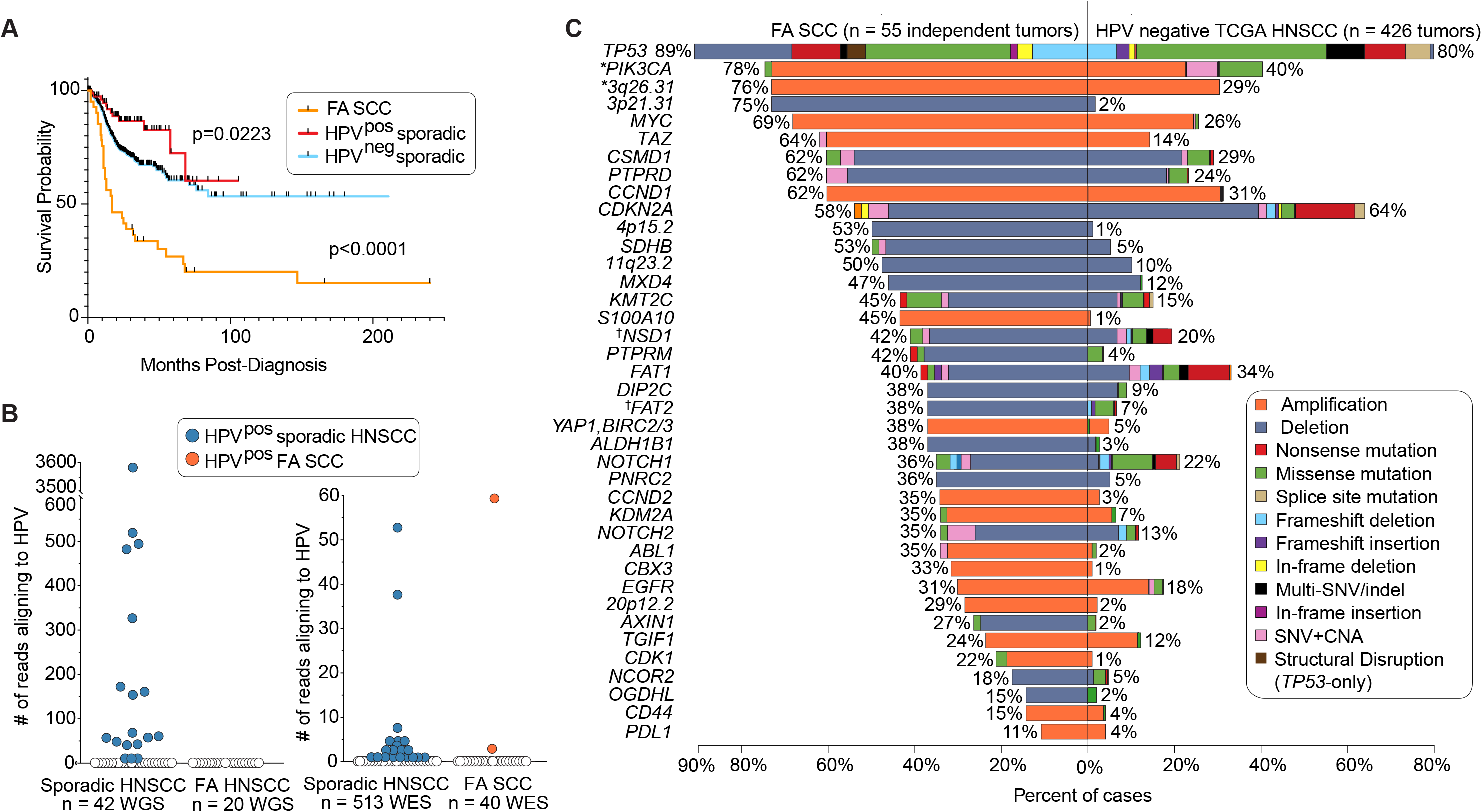
Comparison of the mutational landscape between squamous cell carcinomas (SCC) from individuals with Fanconi anemia (FA) and sporadic head and neck SCCs (HNSCC). **a** Cancer-specific survival curves for 41 individuals with FA-associated SCC and complete clinical history, 78 HPV-positive sporadic HNSCC cases, and 419 HPV-negative sporadic HNSCC cases from TCGA. p-value indicates log-rank Mantel-Cox test. **b** Number of paired-end reads aligning without clipping to non-repetitive regions of any HPV genome in FA SCCs (20 WGS and 40 WES) and sporadic HNSCC tumors from TCGA (42 WGS and 513 WES). **c** Comparison of gene alteration frequencies between independent FA SCC tumors (n = 55) and HPV-negative sporadic HNSCC tumors (n = 426), with focal somatic copy number alteration (sCNA) peaks defined by *GISTIC2* [read-depth change of log2(sCNA)≥0.9 (amplification) or log2(sCNA)≤-0.9 (deletion) relative to pooled normal region coverage] and with normalization for tumor purity in both cohorts. SNV is single nucleotide variation. *TP53* and *PDL1 s*CNA frequencies were determined manually. * and † indicate genes contained within the same focal sCNA peak.

## Majority of the FA SCCs are HPV-negative, TP53 mutant, and display frequent copy-number alterations

HPV infection is a well-characterized driver of sporadic HNSCC, particularly for younger patients^17, 21^. Although there is evidence that the FA pathway protects against HPV replication, the importance of HPV in FA SCC etiology has been debated^22–26^. To identify HPV sequences integrated within the FA SCCs, whole genome sequences from 20 FA and 42 sporadic HNSCCs (The Cancer Genome Atlas [TCGA]) were aligned against all genotyped HPV strains. No HPV sequences were identified in the FA HNSCCs, while we detected such sequences in 18 HPV positive sporadic HNSCCs **(Fig 1B)**. Using whole exome sequencing of 40 FA SCC tumors across multiple tissue sites, we found only two HPV positive cases both of gynecologic origin **(Fig 1B, Supplemental Table 2)**. Consistent with the low HPV infection incidence, *TP53* mutations were identified in 89% of the FA SCCs **(Fig 1C)**.

With the notable exception of *TP53*, we found a reduced rate of single nucleotide variants (SNV) and small insertion/deletion events (indels) perturbing known SCC driver genes in FA SCCs compared to HPV-negative sporadic HNSCC **(Fig 1C, Extended data Fig 1E**). For example, only one FA SCC sample harbored a PIK3CA-missense mutation (1.8% of samples), as compared to 17% of the HPV-negative sporadic HNSCCs (TCGA). A similar lack of point mutations was seen in *CDKN2A*, *FAT1, NOTCH1*, *NSD1*, and several other tumor suppressors. A global analysis of somatic SNV and indel events uncovered a reduced exonic mutation rate relative to sporadic HNSCC **(Extended data Fig 2A, Supplemental Table 3),** most likely due to an earlier age of cancer onset in FA patients. Deconstruction of FA SCC SNV de-novo profiles into Catalogue of Somatic Mutation In Cancer (COSMIC) mutational signatures, revealed the presence of mutational signatures SBS1 & SBS5 (cytosine deamination and unknown etiology), SBS2 & SBS13 (APOBEC), and SBS39 (bulky DNA adducts) **(Extended data Fig 2C**). However, we found no evidence of the general HR-deficiency associated SBS3, or the SBS4 signature associated with cigarette smoke exposure. Analysis of the de-novo indel mutation profiles from FA SCCs revealed signatures ID6 (MMEJ/NHEJ), ID8 (NHEJ), ID10 (large insertions at tandem repeats), and ID4 (small deletions at tandem repeats) **(Extended data Fig 2D)**. ID10, ID4, and ID6 were not enriched in sporadic HNSCC.

Instead of a high SNV/indel load, we find that chromosomal instability, a hallmark of FA patient cells, is the major force driving FA SCCs^1–3^. Analysis of high-level, focal somatic copy-number alteration (CNA) in 55 independent FA SCCs revealed a significantly elevated somatic CNA frequency relative to HPV-negative sporadic HNSCCs **(Fig 1C, Extended data Fig 1E, Fig 3 and Fig 4)**. Among the amplified loci are those harboring *PIK3CA* (amplified in 76% of samples), *MYC* (69%), *TAZ* (62%), *CCND1* (62%), *YAP1/BIRC2/BIRC3* (38%)*, CCND2* (35%), KDM2A (34%), and *EGFR* (31%). Among deleted loci are *PTPRD* (deleted in 62% of samples), *CSMD1* (58%), *CDKN2A* (55%), *SDHB* (51%), *MXD4* (47%), *NSD1/FAT2* (38%), *KMT2C* (35%), *NOTCH1* (33%), and FAT1 (33%). Overall, CNAs perturbing oncogene and tumor suppressor loci were substantially more frequent in FA SCCs than in sporadic HNSCCs. Strikingly, each FA SCC carried multiple amplifications and deletions across numerous loci **(Extended data Fig 1E, Supplemental Table 4)**. The most significant co-occurrence was between *PIK3CA* and *MYC* amplifications, which was present in 55% of FA SCCs.

## FA SCCs are characterized by an elevated burden of structural variants enriched for deletions and localizing to fragile sites

Using whole genome sequencing (WGS) datasets, we next compared the somatic structural variant (SV) landscape of FA tumors (20 SCC and 2 adenocarcinomas) to that of sporadic HPV-negative (n=24) and HPV-positive (n=18) HNSCCs, as well as breast and ovarian cancers driven by *BRCA2* (*BRCA2^mut^*, n=41) or *BRCA1* (*BRCA1^mut^*, n=24) mutations **(Supplemental Table 5)**. FA tumors displayed a median 2.2-fold increase in the total SV burden compared to HPV-negative HNSCCs **(Fig 2A)**. This difference remained unchanged after exclusion of adenocarcinoma samples. FA tumors had, on average, 45% more SVs than *BRCA2^mut^* carcinomas, but 30% fewer SVs than *BRCA1^mut^* carcinomas, which are highly enriched for tandem duplications (TDs) **(Fig 2A)**^27–29^. FA SCCs exhibited a significantly increased burden of SVs across all classes with a 2.2-fold increase in deletions, 3.5-fold increase in translocations, 2-fold increase in inversions, and 1.5-fold increase in TDs when compared to HPV-negative sporadic HNSCC **(Fig 2B)**. Deletions were significantly enriched in FA SCCs **(Fig 2C)**. While SV breakpoints in FA SCC clustered at oncogenic and tumor suppressor loci, breakpoints were also distributed at low frequencies throughout the genome **(Extended data Fig 5),** suggestive of a model whereby continuous DSB generation due to abnormal DNA repair provides the substrate of genetic variation within the pool of epithelial cells for selection to act upon.

**Fig. 2.**
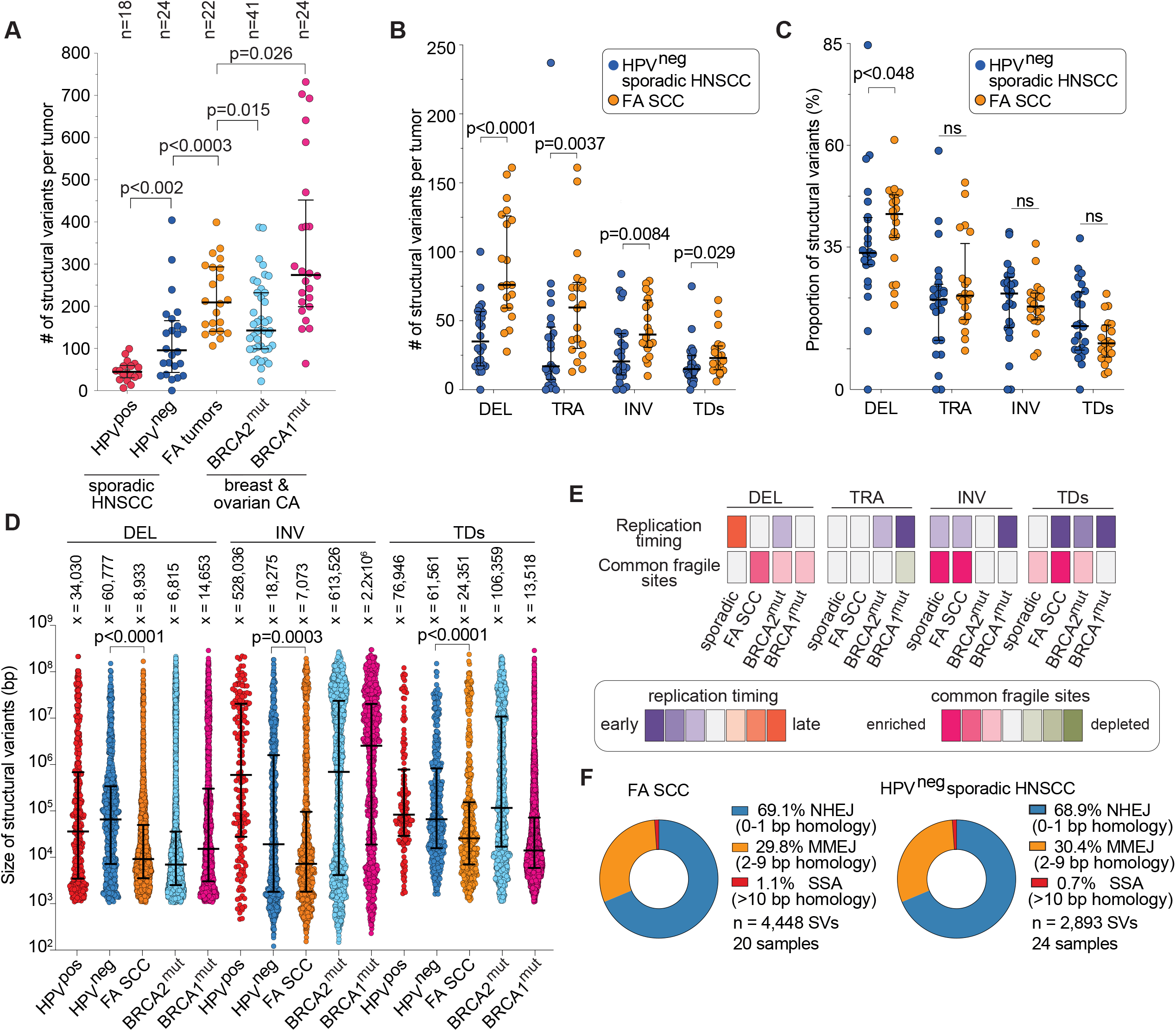
Analysis of the structural variant landscape in SCCs from FA patients. **a** Comparison of somatic structurial variant (SV) numbers in the whole genomes of FA-associated cancers (20 SCC and 2 adenocarcinomas), HPV-positive sporadic HNSCC, HPV-negative sporadic HNSCC, BRCA2-deficient (BRCA2^mut^), or BRCA1-deficient (BRCA1^mut^) breast and ovarian tumors. **b** SV counts in FA SCC (n=20) and HPV-negative sporadic HNSCC (n=24) cohorts categorized by class: deletion (DEL), translocation (TRA), inversion (INV), and tandem duplication (TDs). INV include reciprocal inversions, fold-back inversions and complex intrachromosomal rearrangements with inverted orientation **c** Data in b showing proportion of all SVs represented by each SV class. **d** Size distribution of SV classes in the indicated tumor cohorts. Size (in base pairs; bp) is defined by intrachromosomal distance between left and right breakpoints **e** Replication timing and common fragile site localization of SV breakpoints, stratified by both SV class and tumor cohort. Colour scale indicates strength of correlation. Detailed data is shown in Extended data Figure 6. **f** Mechanism of breakpoint resolution in FA SCC (n=20) and HPV-negative sporadic HNSCC (n=24) cohorts, categorized by the double-strand DNA break repair pathways: non-homologous end joining (NHEJ), microhomology-mediated end joining (MMEJ), and single strand annealing (SSA). Indicated is the proportion (%) of breakpoints predicted to have been repaired by each pathway, based on previously established homology parameters. Mann-Whitney U test two-tailed exact p-values are indicated, with median and IQR shown (**a-d**).

The SV size (length of DNA between two intrachromosomal breakpoints) predominantly ranged between 1-100kb in FA SCC **(Fig 2D)**. HPV-negative sporadic HNSCCs displayed a clustering of mid-size deletions with a median of 60kb (interquartile range [IQR] of 6-291kb), while FA SCCs exhibited a clustering of small deletions with a median size of 9kb (IQR 3-42kb). Sizes of deletions were similar between FA SCC and *BRCA2^mut^* samples. FA SCC inversions were also small and clustered with a median size of 7kb (IQR 2-90kb). This differed from the more evenly spread inversions of HPV-negative HNSCC with a median of 18kb (IQR 2kb-1.4Mbp), the bimodal clustering of small and large inversions in *BRCA2^mu^*^t^, and the large inversion cluster of *BRCA1^mut^* samples. Like deletions and inversions, TDs in FA SCCs were also small, at 24kb (IQR 7-140kb), in comparison to HPV-negative HNSCC and *BRCA2^mut^* samples, with median sizes of 61kb (IQR 14-632kb), and 106kb (IQR 9kb-5.7Mb) respectively. FA SCCs did not share the characteristic TD enrichment observed in *BRCA1^mut^* tumors; however, the TD size distribution was similar between these two cohorts.

Genomic replication timing has been demonstrated to have a strong association with SV formation. Deletions are enriched in late-replicating regions, while TDs and unbalanced translocations occur preferentially in early-replicating regions across the Pan-Cancer Analysis of Whole Genomes^30^. We assessed replication timing and SV localization in FA SCC in comparison to sporadic HNSCC, *BRCA2^mut^*, and *BRCA1^mut^* cancers. Compared to reference timing profiles^31^, we found that TD breakpoints in FA SCC, *BRCA1^mut^*, and *BRCA2^mu^*^t^ tumors, but not sporadic HNSCC, are highly associated with regions of early genomic replication **(Fig 2E, Extended data Fig 6C)**. Deletions, on the other hand, were found in late-replicating regions only in sporadic HNSCCs. Consistent with the FA pathway being important at common fragile sites^32^, we found that deletion, inversion, and TD breakpoints were greatly enriched at these loci in FA SCCs. Only inversions, and to lesser extent TDs, were enriched at common fragile sites in sporadic HNSCCs **(Extended data Fig 6A)**. TD breakpoints in FA SCCs and *BRCA1^mut^*, and translocations in *BRCA1^mut^,* correlated with early replication fragile sites **(Extended data Fig 6B)**.

FA pathway-deficient cells have been demonstrated to harbor rearrangements driven by microhomology-mediated end joining (MMEJ)^7^. We assessed the frequency of predicted non-homologous end joining (NHEJ) [0-1bp breakpoint homology (BH)], MMEJ [2-9bp BH], and single-strand annealing (SSA) [>10bp BH] and found that FA pathway status does not alter the mechanism of DNA double strand break resolution **(Fig 2F)**.

## SVs in FA SCCs frequently form complex chains comprised of unbalanced translocations, fold-back inversions, and templated insertions, contributing to oncogene amplifications

FA pathway-deficient cells are known to contain complex radial chromosomes that can be visualized on metaphase spreads^1, 2^. We hypothesized that cycles of their formation and breakage during tumorigenesis leads to the development of rearrangement chains. To assess their presence, we performed 10X linked-read WGS on four FA SCC samples, which allowed for phasing of structural events across long genomic segments. We observed that SVs frequently occurred in long chains forming complex chromosomal rearrangements **(Fig 3A and B)**. Samples had between 23 and 32 unique chains with a mean of 4.6 SVs per chain **(Extended data Fig 7A and B)**. The largest observed chain contained 43 SVs. These chains were enriched for duplications, translocations, and deletions, and frequently localized to oncogene-containing regions of chromosomes 3, 7, 8, 9, and 11 **(Extended data Fig 7C and D)**.

**Fig. 3.**
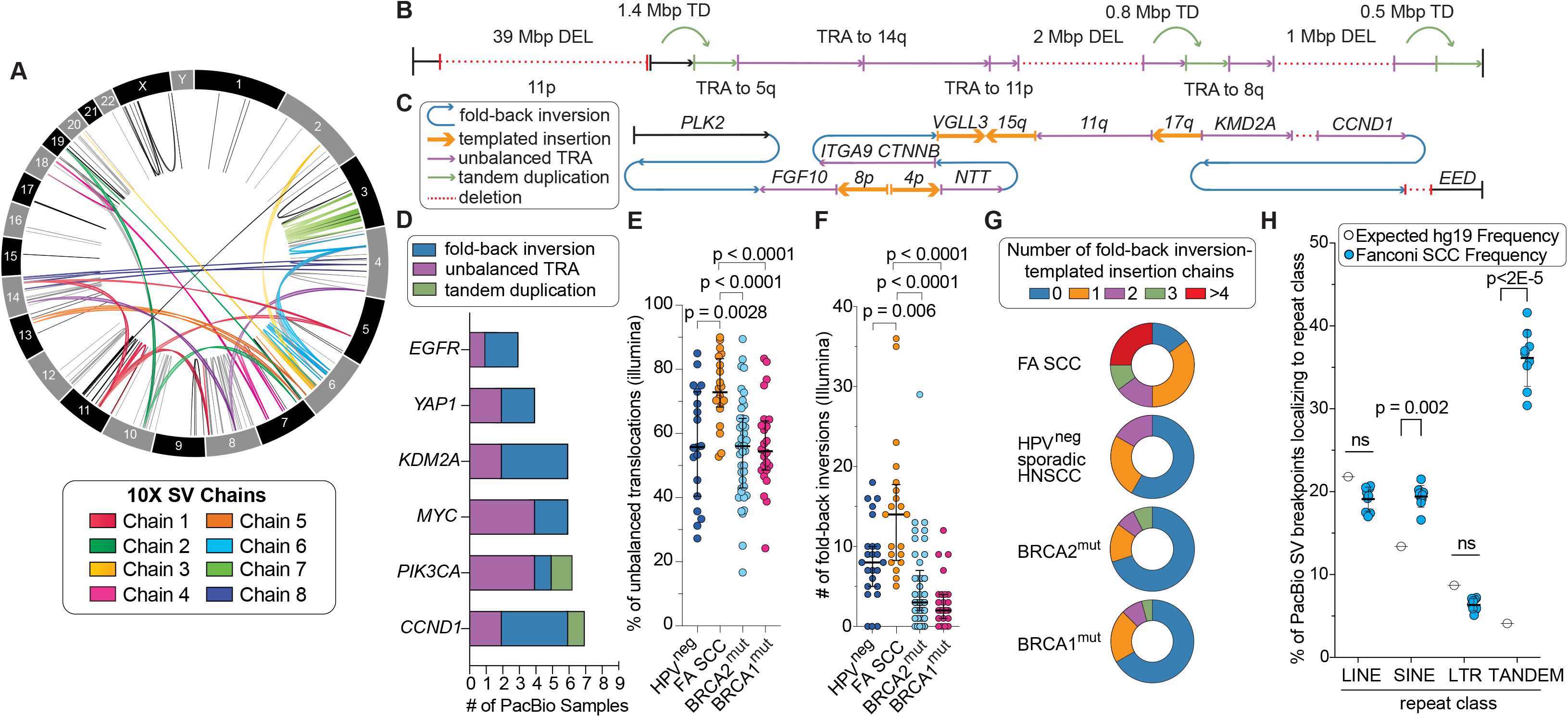
Analysis of complex FA SCC structural variants identified by 10x linked-read, PacBio long read, and Illumina WGS. **a** Circos plot of somatic structural variants (SVs) larger than 30kb detected using 10X linked-read WGS in FA SCC (sample F17P1). 8 selected multi-SV chains are highlighted using distinct colors, with the outer ring segmented by chromosome number. A chain is defined as a minimum of 4 barcode-linked breakpoints (≥ 2 SVs). **b** Illustration of SV chain #1 from panel a. Color legend is shared with panel c. Arrows indicate orientation of each segment relative to hg19 reference genome. **c** Illustration of a somatic SV chain containing unbalanced translocations, fold-back inversions, and templated insertion chains present in FA SCC sample F45P1. **d** Deduced amplification mechanism at select oncogenes in FA SCC as assessed by PacBio sequencing. **e** Proportion of translocations events that are unbalanced (non-reciprocal and copy-number altering) among FA SCC (n=20), HPV-negative sporadic HNSCC (n=19), BRCA2^mut^ carcinomas (n=41), and BRCA1^mut^ carcinomas (n=24). 5 sporadic HNSCC samples with ≤3 translocation events were excluded. **f** Number of fold-back inversion events in the same cohorts. **g** % of the samples in each cohort with 0, 1, 2, 3, or more than 3 unique FBI-TIC chains. **h** Comparison of expected (hg19 reference) vs. observed percentage of somatic SV breakpoints localizing to indicated repeat class. Breakpoints from 9 FA SCC PacBio samples were used. Mann-Whitney U test two-tailed exact p-values are indicated, with median and IQR shown **(e-g).**

To capture complex SV subtypes, we used the PacBio long-read platform to sequence whole genomes of 9 FA SCC samples. While the increased sensitivity of long-reads increased the number of captured SVs, the PacBio data displayed a similar enrichment for somatic deletions as Illumina WGS data **(Extended data Fig 7E and F)**. PacBio sequencing revealed a high frequency of unbalanced translocations (UBT) and fold-back inversions (FBI) that were frequently connected to form complex rearrangements involving multiple oncogenes **(Fig 3C)**. UBT and FBI events drove sharp copy number amplification at key oncogenes including *CCND1*, *PIK3CA, MYC, KDM2A, YAP1-BIRC2/3,* and *EGFR* **(Fig 3D, Extended data Fig 7H).** UBT and FBI segments were often bridged by one or more templated insertions of 1 kbp or less, copied from distant loci **(Fig 3C).** When strung together, these short insertions formed templated insertion chains (TIC)^30^, which connected multiple intrachromosomal and interchromosomal loci. The analysis of PacBio data prompted us to quantify the frequency of unbalanced translocations **(Figure 3E and Extended data Figure 7I)**, FBIs **(Figure 3F)**, and FBI-TICs **(Figure 3G)** present in FA SCC, HPV-negative HNSCC, BRCA2^mut^, and BRCA1^mut^ cohorts **(Supplementary Tables 5 and 6).** We found that all three classes of SVs were significantly enriched in FA SCC relative to the other cohorts, demonstrating that the FA repair pathway protects against such events that can drive oncogenic amplification.

Repeat-rich loci are susceptible to higher levels of replication stress^33^. Using PacBio data, which is able to accurately align these loci, we found that somatic SV breakpoints in FA SCCs preferentially localized to repeat regions, relative to the expected background of the hg19 reference genome (83% observed vs 51% expected) **(Extended data Figure 7G)**. Breakpoints were 8-fold enriched in simple/tandem repeat regions (regions of 1-5bp repeat patterns), and 1.5-fold enriched in SINE elements, but were not enriched in LINE elements or LTRs **(Fig 3H)**. Regions of SV breakpoints (+/−100 bp around the breakpoint) were mildly enriched in GC content with a median +7% shift relative to the hg19 GC-background **(Extended data Fig 7L)**. While detection capacity is limited with Illumina short-read data in repetitive regions, especially at tandem repeats, we found enrichment of somatic SV breakpoints within tandem and SINE elements when comparing FA SCC vs. sporadic HNSCC **(Extended data Fig 7K)**. However, we found that retrotransposition events were not increased in FA SCC relative to HPV-negative HNSCC **(Extended data Figure 7J)**.

To correlate the increased genomic instability of FA SCCs with transcriptional output, we compared the transcriptomic landscape of FA SCCs against sporadic HNSCCs **(Supplementary Data Table 7).** We performed RNAseq on six FA SCC tumors and compared expression data with that of sporadic HNSCC. Although data were limited by the number of available FA SCC samples, transcriptional expression mirrored DNA sequencing findings, with many of the amplified genes being expressed at higher levels and deleted genes being expressed at lower levels in FA samples relative to sporadic HNSCCs **(Extended data Fig 8A and D).** We additionally found that the global DNA damage response was upregulated in FA SCCs and aldehyde detoxification enzymes were downregulated compared to sporadic HNSCCs **(Extended data Fig 8B).** Gene expression of *ALDH/ADH* genes associated with acetaldehyde, retinaldehyde, and lipid aldehyde processing was lower in FA SCCs **(Extended data Fig 8C)**. We also observed transcriptional downregulation of the MHC Class I antigen processing and presentation pathway **(Extended data Fig 8E).** Comparison of methylation patterns between FA SCCs and sporadic HNSCC using EPIC 850K methylation arrays revealed no significant differences between the two cohorts **(Supplementary Data Table 8)**.

Somatic mutation and hypermethylation of FA pathway genes has been proposed to occur in a subset of sporadic HNSCCs^34–36^ although the functional consequences of the identified genomic changes have not been assessed. Re-analysis of the TCGA HNSCC data revealed that somatic deletions of *MAD2L2* (*FANCV*), *ALDH2, RAD51* (*FANCR*), and *XRCC2* (*FANCU*) correlated with a highly copy-unstable subset of tumors, representing close to 13% of HPV-negative sporadic HNSCCs **(Extended Data Fig 9A)**. This subset displayed significantly increased CNA frequencies at HNSCC driver loci relative to the complete HPV-negative HNSCC cohort **(Extended Data Fig 9B)**. We note, however, that the deletion of these genes may not have driven the observed instability but may simply represent passenger mutations occurring in tumors with high levels of genomic instability.

Acquisition of high copy number variation in a subset of sporadic HNSCCs may alternatively be explained by the functional overload of a genetically unaltered FA repair pathway by exogenous aldehydes. Most notable of these are acetaldehyde, a byproduct of ethanol metabolism, as well as formaldehyde and acrolein present in tobacco smoke. To determine whether exposure to exogenous DNA crosslinking agents correlated with level of CNA formation, we stratified HPV-negative sporadic HNSCC tumors by patient pack-year smoking history **(Extended Data Fig 9C)**. We found that the median number of focal CNAs was elevated in patients who smoked relative to non-smokers, and that this difference widened as the number of pack-years increased.

Subsequently, we ranked sporadic HPV-negative HNSCC tumors by level of focal copy-number instability and found that the most unstable quartile demonstrated a strong enrichment for acetaldehyde (DBS2), smoking carcinogen (SBS4 and ID3), and NHEJ (ID8) mutational signatures relative to the most copy-stable tumor quartile. We further found that tumors from smokers in the most unstable quartile had the highest exposure to tobacco when compared with those in the most stable quartile **(Extended data Fig 9D-G)**. We propose that high aldehyde exposure in this subset of tumors during the early stages of tumorigenesis generates extensive ICL damage, which may overwhelm a genetically unaltered FA DNA repair pathway – particularly in the setting of mutated TP53. This may lead to a high CNA rate that fuel further carcinogenesis (see discussion).

## A mouse model of FA SCC displays accelerated growth, swift epithelial-to-mesenchymal transition, and strong induction of autonomous inflammatory signaling

To test if a combination of FA pathway and TP53 tumor suppressor deficiency may recapitulate the increased SV load and hasten SCC development, as seen in human FA tumors, we created a mouse serial allograft model of FA SCC. *Fanca^−/−^ Trp53^−/−^ and Fanca^+/+^ Trp53^−/−^* neonatal keratinocytes were immortalized and transformed by overexpression of *Ccnd1* and *HRAS^G12V^* respectively. These keratinocytes were serially engrafted in Nude mice **(Fig 4A)**. During the 11 cycles of engraftments, we quantified tumor growth and performed RNAseq, Illumina WGS, histopathology, and protein expression analysis. This experiment revealed that *Fanca^−/−^* keratinocytes exhibit dramatically accelerated tumor growth compared to their *Fanca^+/+^* counterpart. The difference became apparent at the first engraftment cycle **(Fig 4B and C)**, despite pre-engraftment *Fanca^−/−^* and *Fanca^+/+^* keratinocytes having identical growth *in vitro* **(Extended data Fig 10A)**. Growth acceleration was observed through multiple subsequent engraftments, but the difference between *Fanca^−/−^* and *Fanca^+/+^* dissipated by the time of the 11^th^ engraftment **(Extended data Fig 10B)**.

**Fig. 4.**
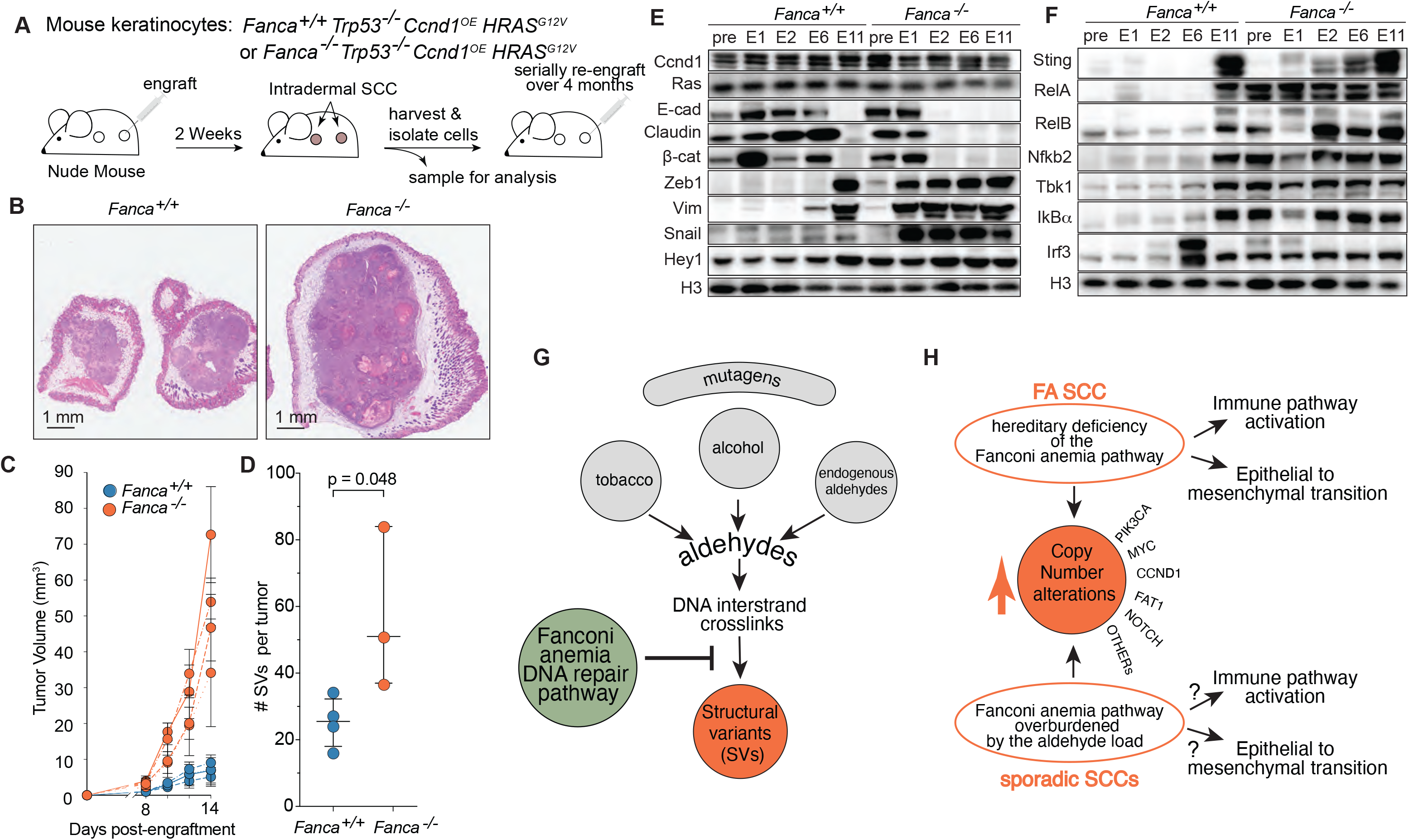
Characterization of a murine FA SCC model. **a** Schematic showing the serial engraftment of murine keratinocytes of the indicated genotypes. **b** Representative micrographs of H&E-stained tumor sections derived from *Fanca^+/+^* and *Fanca^−/−^* keratinocytes at the end of the first engraftment cycle. **c** Tumor volumes during first engraftment of *Fanca^+/+^* and *Fanca^−/−^* keratinocytes. Each point represents the mean volume of four tumors from one mouse with standard error shown. Four replicates, each comprised of four tumors engrafted on a single mouse, are indicated by four separate curves. **d** Somatic SV event counts in 3 *Fanca^−/−^* and 4 *Fanca^+/+^* independently grown tumor replicates at the 6^th^ engraftment cycle (∼2 months). Bars indicate median-IQ range. Two-tailed, unpaired t-test p-value is shown. **e** and **f** Protein levels of epithelial and mesenchymal (e) and inflammatory (f) markers measured by western-blotting across engraftment time points for *Fanca^+/+^* and *Fanca^−/−^* genotypes. See Extended Data Fig 10F for transcript-level expression of these and additional markers. **g** Fanconi anemia pathway protects against copy number alterations driving squamous cell carcinoma development. By removal of DNA interstrand crosslinks created by endogenous and exogenous aldehydes, the FA pathway functions to prevent formation of structural variants including deletions, unbalanced translocations, and foldback inversions **h** In FA patients, hereditary deficiency of DNA interstrand crosslink repair leads to formation of structural variants, which in the setting of TP53 deficiency leads to high copy number alteration in key oncogenes and tumor suppressors. We propose that the appearance of copy number alterations in sporadic HNSCC may result from the functional overload of a genetically unaltered FA pathway by exogenous aldehydes in tobacco and alcohol, especially in the setting of TP53 mutation when the DNA repair pathways cannot be properly induced. Immune pathway activation and epithelial to mesenchymal transition are additional features of FA SCC that contribute to aggressive nature of these tumors.

WGS of the mouse tumors revealed a median two-fold increase of somatic SVs in the tumors derived from *Fanca^−/−^* relative to *Fanca^+/+^* keratinocytes at the 6^th^ cycle **(Fig 4D)**, with the largest increase resulting from inversions **(Extended data Fig 10C and D)**. It is unclear whether this is due to species-specific differences in DNA repair, the genetic background of the keratinocytes, or the influence of *HRAS^G12V^*. While pre-engraftment *Fanca^−/−^* and *Fanca^+/+^* keratinocytes exhibited similar gene expression profiles, the first engraftment induced rapid transcriptomic changes exclusively in *Fanca^−/−^* tumors. *Fanca^−/−^* keratinocytes invoked expression programs characteristic of epithelial-to-mesenchymal transition (EMT), intracellular inflammatory signaling, TGF*β* pathway activation, cancer stem cell transition, and cellular metastasis, among several others **(Extended data Fig 10E and F, Supplementary Table 9)**. EMT marker induction at the first engraftment cycle was confirmed at the protein level, with a strong rise in SNAIL and ZEB1 levels accompanying the appearance of Vimentin. Subsequent disappearance of epithelial markers including E-cadherin, Claudin, and *β*-catenin occurred at the second engraftment **(Fig 4E)**. *Fanca^+/+^* keratinocytes eventually underwent EMT by the 11^th^ engraftment cycle, and this correlated with a narrowing of the growth rate difference between *Fanca^+/+^ and Fanca^−/−^* cells **(Extended data Fig 10B)**. Protein expression studies also revealed strong activation of intracellular inflammatory pathways, including canonical (RELA, TBK1, IRF3, IKBA) and non-canonical (RELB, NFKB2) NFkb pathways, already present in the pre-engraftment *Fanca^−/−^*, but not *Fanca^+/+^* keratinocytes **(Fig 4F)**. Over the course of multiple engraftment cycles, we observed significant upregulation of the dsDNA-sensor *STING* in *Fanca−/−* tumors **(Fig 4F and Extended data Fig 10F)**. STING is activated by cGAS stimulation at micronuclei formed in the setting of genome instability, including in primary cells from FA patients^37–40^. Although these phenomena need to be further studied, our experiments suggest that the robust induction of EMT occurring in the setting of high DNA damage and immune pathway activation may contribute to the aggressive nature of the FA-associated SCCs. Collectively, the effects of these changes go beyond the influence of the FA pathway deficiency on structural and copy number variation in oncogenes and tumor suppressors, as they occur so swiftly in the mouse allograft model.

## Discussion

Sporadic HNSCCs and esophageal SCCs are rich in somatic CNAs affecting multiple tumorigenesis-associated genes^21, 41, 42^. Analyses presented here demonstrate that loss of the FA repair pathway increases CNA frequency in tumors, through induction of simple and complex structural rearrangements **(Fig 4G).** This phenotype is consistent with a lack of proper DNA repair during replication, leading to unregulated formation of multiple DNA double strand breaks that are repaired by NHEJ and MMEJ pathways. The breakpoints of SVs in FA SCCs were enriched at common fragile sites and other potentially difficult to replicate regions, consistent with known function of the FA pathway at those sites. High numbers of fold-back inversions, unbalanced translocations, and deletions present in the FA SCCs point to mechanisms of inappropriate processing of stalled replication forks in the setting of FA repair pathway deficiency, leading to genomic copy number alterations.

Passenger mutations are clearly present among the focal amplification and deletion peaks identified in FA SCCs. However, the high frequency of genomic alterations in FA tumors provides a rich substrate pool for selection to act upon during tumor progression. This results in recurrent genomic events affecting known sporadic HNSCC driver genes, including *PIK3CA, MYC, YAP1, CCND1, CSMD1, PTPRD, MXD4*, *EGFR,* and others. The occurrence and co-occurrence of these genetic lesions in FA SCC is substantially higher than in sporadic HNSCCs, which may partially explain the early-onset and aggressive nature of FA SCCs. Analysis of the increased SV load in FA SCCs is an opportunity to nominate additional SCC tumor suppressors and oncogenes that have been overlooked in the analysis of sporadic cancers. Functional studies will be necessary to assess the role of such uncharacterized genes to determine if they contribute to the progression from normal keratinocyte to invasive SCC.

A striking finding in this study of FA SCCs is that in the sea of structural variants, *TP53* is the only gene that is significantly inactivated by SNVs and indels. *TP53* inactivation is an early event in the pathogenesis of sporadic HNSCC and likely occurs early in the development of FA-associated tumors. This permits rearrangements to develop and to be selected for, without induction of apoptosis or senescence. Presence of *TP53* mutations should be validated in FA patients as an early tumor marker and a potential screening biomarker for future cancer prevention studies.

The epidemiology of sporadic aerodigestive tract SCCs, including HNSCCs and esophageal SCCs not associated with HPV, points to an intimate connection between tumorigenesis and mutagen exposure, particularly, tobacco, alcohol consumption, and environmental pollution^17^. Although resulting SNV and indel mutations contribute to tumorigenesis in sporadic HNSCCs, notably through mutagenesis of *TP53*, *CDKN2A*, and *PIK3CA*, we propose that inefficiently cleared ICL damage from these mutagens additionally contributes to genome instability in sporadic HNSCCs through induction of structural variation. Multiple lines of evidence lead us to this conclusion. First, the presence of *ALDH2* mutations predisposes to HNSCCs in the general population. Second, there exist a strong genetic interaction between aldehyde metabolism and the FA pathway. Finally, a clear correlation is present between tobacco smoke exposure and copy-number instability in sporadic HNSCC. We discuss these in turn.

*ALDH2* mutations in East Asian populations, which result in deficient aldehyde clearance, greatly predispose to HNSCC^43^. Endogenously produced aldehydes such as 4-hydroxy-2-nonenal (HNE), formaldehyde, and malondialdehyde, as well as acetaldehyde (a byproduct of ethanol) are all detoxified by ALDH2. These aldehydes form DNA interstrand crosslinks requiring the FA pathway for repair. This is most clearly illustrated by genetic studies showing that bone marrow failure and leukemogenesis is enhanced by co-mutation of *ALDH2* and the FA pathway, both in mice and humans^7, 8, 44^. The reduced aldehyde clearance results in increased dependency on the FA pathway for genome stability. One could conclude that this interaction is only important in the setting of FA pathway deficiency. However, a recently identified digenic syndrome of ALDH2 and ADH5 deficiency was shown to be associated with an increase in formaldehyde-induced DNA crosslinks^18, 19^. The result is an FA-like disorder in patients, including bone marrow failure and AML induction. Remarkably, these phenotypes were present without a direct mutation of the FA pathway, which clearly demonstrates that even a functional FA pathway can still be overwhelmed by an increased aldehyde load^20^.

Analysis of sporadic HNSCCs performed in this study suggests that the most copy-number unstable tumors were enriched for acetylaldehyde and smoking mutagen exposure signatures. We further found that cigarette smoke exposure is correlated with increased copy-number instability. Thus, we speculate that high genome instability characterized by frequent amplifications and deletions seen in these sporadic HNSCCs may be due to a functionally overwhelmed FA pathway by the elevated aldehyde load derived from tobacco smoke inhalation, alcohol consumption, and environmental pollutants **(Fig 4H)**. The resultant genomic instability in the form of SVs from an overburdened FA pathway is superimposed over known SNV and indel-inducing mutagenesis associated with tobacco use and alcohol consumption.

Our mouse model points to other hallmarks of SCC that are likely driven by DNA damage when the FA pathway is absent or overwhelmed. Most notably, we observed accelerated epithelial-to-mesenchymal transition and increased autonomous inflammatory signaling. The exact mechanism of these transitions remains to be determined but may involve triggering of STING-mediated NFkB pathway activation by DNA damage. In particular, the non-canonical NFkB pathway has been demonstrated to drive EMT in chromosomally unstable cancers^45^. Study of these pathways will be important for understanding disease pathogenesis and improving the treatment of sporadic and FA SCCs. Additional mouse and human tumor models with FA pathway deficiency need to be developed to achieve those goals.

The data presented here indicate a challenging path forward in the development of new therapeutic treatments for FA patients with SCC. While surgical resection remains the first-line treatment, a multi-pronged pharmacological approach might be required to push these cancers into remission. Of the potential targets, *PIK3CA* is the most highly penetrant amplification in 76% of FA SCCs. Nonsteroidal anti-inflammatory drugs and the p110*α*-selective kinase inhibitor *Alpelisib,* recently approved for *PIK3CA*-mutant breast carcinomas, need to be tested as potential preventive and therapeutic agents in FA SCCs^46–48^. *MYC* amplification, present in 69% of samples, is the next obvious target for therapeutic inhibition. While MYC is difficult to inhibit pharmacologically, several experimental inhibitors like *MYCi975* should be tested in preclinical models^49^. With 55% of FA-SCC cases exhibiting strong co-amplifications of *PIK3CA* and *MYC,* combination therapies against these two targets need to be assessed. The EGFR-blocking mAb *Cetuximab* is clinically available^50^, and with 31% of FA SCCs exhibiting *EGFR* amplifications, it needs to be systematically tested in FA preclinical models and FA patients—potentially in combination with low doses of radiation therapy. High deletion frequency of less-explored tumor suppressors like *RASSF1 (3p21.31), PTPRD, CSMD1, MDX4, SDHB, KMT2C, NSD1, FAT1, and NOTCH1* necessitate further exploration and screening for drug development. *PTPRD* mutations, for example, have been associated with increased STAT3 activation, pointing to STAT3 inhibition as a possible treatment option^51^.

The potential of accelerated EMT and hyperinflammation opens another avenue for suppressing the invasive nature of FA SCCs. Pharmacological advances have been made in this area, largely with respect to precise metabolic targeting of EMT cell fate transition^52^ as well as small-molecule inhibitors of STING^53^. Further study is also required to assess the efficacy of checkpoint immunotherapy for FA patients with SCC, which is in use for sporadic HNSCCs^54^. FA patients who have received hematopoietic stem cell transplant have a normally-functioning, but non-native immune system, while those without transplant may have a poorly functioning immune system. It is unclear how well either of the groups will respond to PDL1 and CTLA4 inhibitors.

Finally, to aid clinicians in the selection of appropriate treatment, particularly considering the high relapse rate of FA SCC, our work demonstrates the urgent need for a comprehensive tumor mutation diagnostics pipeline for FA patients. In addition, further development of accurate preclinical models for testing nascent and novel therapies is necessary to improve patient treatment and survival.

## Acknowledgements

We are grateful to participants and their families who donated their tissues to the International Fanconi Anemia Registry (IFAR). We thank the physicians who provided research samples and clinical information and the staff of Fanconi Anemia Research Fund, especially Suzanne Planck for referrals to IFAR. National Disease Research Interchange (NDRI) is acknowledged for providing samples. We appreciate advice from all members of the Laboratory of Genome Maintenance. Staff of the Genomic, Reference Genome, Bioinformatics, and Flow Cytometry resource centers at the Rockefeller University are acknowledged for their expert advice and contribution. Genomic data from non FA SCCs are in part based upon data generated by the TCGA Research Network: https://www.cancer.gov/tcga. This study was supported by Pershing Square Sohn Prize for Young Investigators in Cancer Research (AS), Fanconi Anemia Research Fund (RD, EV, AS), V Foundation grant T2019-013 (AS), National Institutes of Health (NIH) National Heart Lung and Blood Institute (R01 HL120922) (AS), National Cancer Institute (R01 CA204127) (AS), National Center for Advancing Translational Sciences (UL1 TR001866) (RV and AS), NIH award 1DP2-GM123495 (AK). SCC acknowledges support from the Intramural Research Program of the NIH National Human Genome Research Institute. MAS is supported by a Rubicon fellowship from NWO (019.153LW.038) and a KWF Kankerbestrijding young investigator grant (12797/2019-2, Bas Mulder Award; Dutch Cancer Foundation). AS is a Howard Hughes Faculty Scholar.

## Author contributions

AW, FL, RW, AG, SS performed wet lab experiments. AW, MS, FP, RW, AG, ME, FD, KH, TC, HT, JR, AK, MI, SC, PC, AS performed analysis of the data. KP, RN, FL, MJ, TH, JK, OR, AG, AR, AA, BS, DK, AS collected samples and clinical information. RD, EV, JR, SW, JS, GB, MM, JW, MC, FB, BS, DK provided samples and clinical information. OF performed PacBio sequencing. TS provided pathology expertise. RV provided biostatistical expertise. AS and PC supervised the study. AS and SC obtained funding. AW and AS wrote the manuscript with essential input from other authors.

## Additional information

Supplementary data associated with this article contains 9 Supplemental Tables

Supplemental Table 1. Clinical and pathology data for human samples sequenced in this study.

Supplemental Table 2. Alignment of FA SCC and sporadic HNSCC samples against HPV genomes.

Supplemental Table 3. Somatic SNV and Indel calls in FA SCCs and sporadic HPV-negative HNSCCs.

Supplemental Table 4. CNV Calls and Defined Focal amplification and deletion peaks in FA SCC and sporadic HNSCCs.

Supplemental Table 5. SV Calls in FA SCCs, sporadic HNSCCs, BRCA2^mut^, and BRCA1^mut^ tumors.

Supplemental Table 6. Quantification of fold back inversions, fold back inversion-templated insertion chains, retrotransposon element insertions, and select reciprocal inversions (classical inversion) identified in FA SCC and sporadic HNSCC cohort (Illumina WGS).

Supplemental Table 7. FA SCC vs. sporadic HNSCC (TCGA) differentially expressed genes.

Supplemental Table 8. Methylation Array analysis of FA SCCs

Supplemental Table 9. Mouse Tumor SV & RNAseq expression data analysis

## Methods

### Human samples

The International Fanconi Anemia Registry (IFAR) was initiated in 1982 as a prospectively collected database of clinical and genetic information for FA patients^55^. Subjects entered the registry at any point in the disease process with the majority entering at onset of bone marrow failure or cancer occurrence. Patient or next-of-kin consents were obtained and available medical, surgical, pathology, radiation and chemotherapy records were collected from their respective medical centers. Tumor samples were obtained from pathology departments, clinical collaborators and NDRI with proper consents. Most of the normal samples were already available in the IFAR and were collected prospectively. The Institutional Review Board of the Rockefeller University, New York, NY approved this study.

### Distribution of Samples Across Illumina Sequencing Platforms

Illumina whole genome sequencing (WGS) was performed on 22 tumor samples – 19 fresh-frozen (FF) tumors and three tumor-derived primary cell lines. Whole exome sequencing (WES) was completed on 41 Formalin-Fixed Paraffin Embedded (FFPE) tumors. Of the 63 total tumor samples sequenced, 60 were SCC and 3 were adenocarcinomas. 6 SCC samples were WGS/WES pairs (3 tumors sequenced by both WGS and WES) and 4 SCC samples were primary/metastasis pairs (2 primary SCC and 2 associated metastases). Excluding metastasis pairs and samples sequenced using multiple techniques, this yielded 55 independent SCC tumors. Of these independent FA SCCs, 44 were HNSCC, six anogenital SCCs, four esophageal SCCs, and one lung SCC. The adenocarcinomas were from bladder, cervix, and the ampula. Normal tissue was available for 55 of the 63 total samples, predominantly from peripheral blood or primary fibroblasts. See Supplemental Table 1 for details about all sequenced samples.

### Formalin-Fixed Paraffin Embedded Human Tumor Whole-Exome Sequencing

FFPE tumor blocks were sectioned, slide-mounted, and H&E-stained to mark tumor boundaries. Adjacent normal tissue was removed by scalpel. Isolated tumor sections were placed in *Qiagen deparaffinization buffer* at 56°C for 5 minutes. Samples were incubated in Proteinase K and crosslink reversal buffer (*Qiagen Buffer FTB*) for 1 hour at 56°C for protein digestion, followed by 90°C for 1 hour to reverse-crosslink DNA. Uracil-N-glycosylase (UNG) was added to each sample and incubated at 50°C for 1 hour to correct fixation-induced cytosine deamination. Samples were treated with RNAse A, followed by complete lysis (*Qiagen Buffer AL* & ethanol). DNA was captured, washed, and eluted on *Qiagen gDNA* capture columns. DNA was subsequently treated with *NEBNext FFPE Repair Mix* and purified by *Ampure XP beads*. *Agilent Tapestation was used to determine DNA size* and DNA was submitted for whole-exome sequencing. All FFPE tumor samples were paired with patient-matched normal DNA. 25 normal samples were pre-BMT peripheral blood, 10 were primary fibroblasts, and 6 were normal tissue FFPE samples. *Novogene* performed exome capture (*Agilent SureSelect XT V6* & *IDT xGen Exome*) and library preparation. FFPE tumor samples were 2×150bp sequenced to 12Gbp of raw output, and normal samples were sequenced to 6Gbp of raw output.

### Fresh-Frozen Human Tumor Illumina Whole-Genome Sequencing

DNA was extracted from 19 fresh-frozen tumor biopsies and 3 primary tumor lines using the *Qiagen Blood and Cell Culture* extraction kit. Tissue was lysed in Qiagen Buffer GL containing Proteinase K, Qiagen Protease, and RNAse A. DNA was captured on the *Qiagen Genomic Tip* column, washed, and eluted by gravity-flow. *Agilent Fragment Analyzer* was used to check suitability for short-read Illumina WGS and long-read PacBio sequencing. 9 samples with DNA >40kbp were concurrently sequenced using PacBio and Illumina and the remaining 13 tumor samples were sequenced by Illumina WGS only. Patient-matched normal DNA for 15 tumors was extracted from primary fibroblasts or pre-BMT peripheral blood. Illumina WGS sequencing was performed at *New York Genome Centre* (*NYGC*) and *National Institutes of Health (NIH)* using *PCR-free TruSeq* library prep. Tumor samples were sequenced to 60x genome coverage and normal samples were sequenced to 30x genome coverage.

### Human Tumor PacBio Long-Read Whole-Genome Sequencing

6 fresh-frozen tumors and 3 primary tumor cell lines were PacBio sequenced at the *Rockefeller University Reference Genome Resource Center.* PacBio libraries were prepared using the *SMRTbell Express Template v2* library prep kit, using 26G needle-sheared high-molecular weight (HMW) DNA as input. Agarose-plug DNA extraction was performed for the 3 primary tumor lines. 2 samples were sequenced on the *PacBio Sequel 1*, each with 3 x 1M SMRTcells for 10x average genome coverage. 7 samples were sequenced on the *PacBio Sequel 2*, each with an 8M SMRTcell for 30x average genome coverage. 1 sample in this latter cohort yielded lower than-expected output (∼10x genome coverage).

### Human Tumor 10X Linked-Read Whole-Genome Sequencing

4 tumor samples (3 primary tumor lines and 1 fresh-frozen tumor) had sufficiently sized ultra-HMW DNA (median 100kbp+) for *10X linked-read WGS*. Tumor DNA was extracted using the *Circulomics Big DNA* bead kit, using the ultra-HMW elution option. Cell line DNA was extracted using agarose-plug lysis. DNA was run on *Chromium WGS* partitioning chips for GEM encapsulation and DNA fragment barcoding. Each 10X WGS library was sequenced to estimated 60x coverage at the *New York Genome Center*.

### Human Tumor DNA Methylation Analysis

6 FA-SCC tumors and matched normal tissue with sufficient DNA yield were selected for DNA methylation analysis on the *Illumina EPIC Methylation Array* platform. Library preparation and methylation calls were performed at the *NIH Genomics Core*.

### Human Tumor Bulk RNAseq

RNAseq was performed on 6 FA-SCC tumors with sufficient tissue material. Total RNA was extracted using *Trizol* methodology. Purified RNA was processed at *New York Genome Center* with polyA-mRNA capture prior to cDNA synthesis and 2×150bp PE sequencing to >30 million reads/sample.

### Mouse Keratinocyte Line Generation

All mouse studies were approved by the Rockefeller University institutional animal care and use committee. *Fanca^+/−^* 129S mice (*gift of Markus Grompe*) and *Trp53^+/−^* B6/129S mice (*B6.129S2-Trp53tm1Tyj/J* – *Jackson Labs*) were crossbred to produce *Fanca^+/+^ x Tp53^−/−^* and *Fanca^−/−^ x Tp53^−/−^* offspring. At post-natal day 4, a neonate from both genotypes was humanely euthanized and keratinocytes harvested from back-skin tissue via dispase-seperation.^56^ Primary keratinocytes were plated on *3T3-J2* feeder cells until colony formation. On day 5 post-plating, keratinocytes of both genotypes were transduced with *pZIP-mCMV-Ccnd1-IRES-mCherry* vector was applied to immortalize them. Cells were subsequently FACS-sorted for integrin a6^hi^ x integrin b1^hi^ x mCherry^+^ markers, producing basal^57^ (a6^hi^ x b1^hi^) *Ccnd1^OE^* immortalized keratinocyte lines **(extended data Fig 10G)**. *HRAS*^G12V^*-puro* retrovirus *(Addgene plasmid #1768)* was then transduced into the keratinocyte lines and infected cells were puromycin selected, to allow for tumor generation upon engraftment.

### Mouse Tumor Serial Allograft Study

*Fanca^+/+^;Tp53^−/−^;Ccnd1^OE^;HRAS^G12V^ and Fanca^−/−^;Tp53^−/−^;Ccnd1^OE^;HRAS^G12V^* mouse keratinocytes were serially intradermally allografted into backs of *Nude/J* mice *(Jackson Labs)* over 11 engraftment cycles (approx. 17 weeks). Four replicates were created for both *Fanca^+/+^* and *Fanca^−/−^* genotypes, with each replicate carried forward independently through the engraftment cycles. Each replicate consisted of 4 intradermal engraftment sites on the same mouse, which were combined after harvesting at the end of each cycle. At the first engraftment cycle, each engraftment site received 150,000 cells suspended in a 1:1 ratio of DPBS & *Corning Matrigel*. Resulting tumors were grown for a maximum of two weeks or until tumor size reached ∼1.5 cm^3^, whichever occurred sooner. Host mice were humanely euthanized, and tumors were dissected, minced, dissociated, and strained to a single-cell mixture using collagenase and *Trypsin*. Resulting tumor cells were plated at 5% CO_2_ / 3% O_2_ at 37°C, and allowed to recover for 48 hours in low-calcium keratinocyte media *(E-Low Media*)^56^. This ensured consistent cell viability between replicates and permitted *in vitro* analysis of post-transplant cells. The number of engrafted cells was lowered to 100K for both genotypes for engraftments two to five, further lowered to 70K cells for cycles six to nine and reduced to 35K for the 10-11^th^ cycles. One *Fanca^−/−^* replicate was lost at the fifth cycle due to host death, which resulted in removal of this replicate from the sixth engraftment cycle onwards. At the first, second, sixth, and 11^th^ cycles, mice were anaesthetized, and tumors were measured in three dimensions with ISO-calibrated digital calipers (*VWR*) every two days. One repeat first-engraftment cycle experiment for both genotypes was set up to collect tissue for H&E histology of resulting tumors.

### Mouse Tumor Illumina WGS

At the sixth engraftment cycle, *Fanca^−/−^* and *Fanca^+/+^* tumor cells from each genotype replicate were harvested. To remove residual host cells, cells were FACS-sorted for mCherry^+^ directly into cytoplasmic lysis buffer (*Qiagen Buffer C1),* after which the standard *Qiagen Blood and Cell Culture DNA* kit workflow was carried out for DNA extraction. Normal DNA was simultaneously extracted from pre-engraftment *Fanca^+/+^;Tp53^−/−^;Ccnd1^OE^;HRAS^G12V^ and Fanca^−/−^;Tp53^−/−^;Ccnd1^OE^;HRAS^G12V^* keratinocyte lines. Illumina WGS libraries were prepped at *NIH* using the *PCR-free Truseq* protocol and sequenced to 60x genome depth for tumors and 30x genome depth for pre-engraftment normals.

### Mouse Tumor RNAseq

At the first, second, sixth, and 11^th^ engraftment cycles, *Fanca^+/+^;Tp53^−/−^;Ccnd1^OE^;HRAS^G12V^ and Fanca^−/−^;Tp53^−/−^;Ccnd1^OE^;HRAS^G12V^* tumor cells from each replicate were harvested. Cells were FACS-sorted for mCherry^+^ cells directly into RNA-stabilizing cell lysis buffer (*Qiagen Buffer RLT*). Homogenization was performed on *QiaShredder* columns, followed by RNA isolation using the *Qiagen RNAeasy Plus* kit. RNA was simultaneously extracted from pre-engraftment keratinocytes. Total RNA was processed by *Novogene* for polyA+ capture, cDNA synthesis, and 2×150bp PE Illumina sequencing to >30 million reads/sample.

### Histology

Mouse tumor sectioning, H&E staining, and slide scanning was performed at *Histowiz*.

### Western Blotting of Mouse Tumor Cells

*Fanca^+/+^;Tp53^−/−^;Ccnd1^OE^;HRAS^G12V^ and Fanca^−/−^;Tp53^−/−^;Ccnd1^OE^;HRAS^G12V^* mouse tumor cells from the first, second, sixth, and 11^th^ engraftment cycles were scraped from tissue culture plates into DPBS. Cells were lysed in Laemmli buffer containing benzonase and phosphatase inhibitor. Supernatant was quantified by Bradford and DC protein assay (BioRad). 25ug of sample was loaded per well on 8-12% Bis-Tris gels (BioRad) and run for 2.5h/100V. Protein was transferred overnight onto PVDF at 35V/4°C and membranes were blocked in 5% milk. They were incubated with primary antibody overnight at 4°C. After washing, membranes were incubated with secondary HRP-conjugated antibody at RT for 2 hours. They were developed with *Western Lightning ECL Plus* (Perkin Elmer). Blots were imaged on an *Azure c300* chemiluminescent imager.

## Antibodies

Those obtained from Cell Signaling Technology included Ccnd1(E3P5S XP), HRAS (D2H12), Vimentin (D21H3 XP), Claudin-1 (D5H1D XP), *β*-Catenin (D10A8 XP), Snail (C15D3), E-cadherin (24E10), Zeb1 (E2G6Y XP),RelA (p65) (D14E12 XP), Sting (D1V5L),Tbk1 (D1B4), Irf3 (D83B9),RelB (C1E4), IkBa (44D4), Nf-kB2 (p100/p52) (4882), H3 (D1H2 XP). Other antibodies included CD3 (Abcam SP7 ab16669, Hey1 (ProteinTech 19929-1-AP), Integrin a6 (BD Horizon GoH3, BV650), Integrin b1 (BioLegend HMB1-1 APC/Cy7).

## Data Analysis

### Somatic SNV/Indel and Structural Variant Calling

Human WES & WGS sequencing data was processed through the *Wellcome Sanger Institute (WSI) Cancer Genome Project* pipeline^58^. Sequencing data were aligned to NCBI human reference genome (GRCh37) using *BWA-mem*^59^. Duplicate reads were marked by *Picard MarkDuplicates*^60^. Somatic single-nucleotide variants (SNVs) were called with *CaVEMan*^61^ and somatic indels were called by *Pindel*^62^. Post variant calling filters were used to remove artefacts, alignment errors and low-quality variants as described previously in detail^63^. The following filters were applied: (1) common single nucleotide polypmorphisms and artefacts were filtered by the presence in a panel of 75 unmatched normal samples^64^ (2) low-quality variants are filtered by setting the median alignment score threshold (ASMD *≥* 140) and excluding variants for which the majority of reads supporting the variant are clipped (CLPM = 0), (3) counting the number of paired-end reads supporting the variants. Software implementing these filtering steps can be found at https://github.com/MathijsSanders/SangerLCMFiltering. Since many tumor samples suffered from BMT-donor blood contamination, a strict SNV/indel-filtering algorithm was employed next. Variants were deemed somatic if they were either: 1) not present in the *GNOMAD*^65^ germline variant database, or 2) positively reported in the *COSMIC*^66^ database.

Structurial variants (SVs) were called with *BRASS*^67^. SVs were further annotated by AnnotateBRASS (https://github.com/MathijsSanders/AnnotateBRASS) as described previously in detail^63^. In brief, AnnotateBRASS determines per SV: the number of supporting read pairs, the variance in alignment position of read pairs, whether read-pairs are clipped or carry an excess of variants not reported in SNP databases, are in the correct orientation or whether SV-supporting read-pairs are in regions marked by high proportions of other read-pairs aligning to different parts of the genome (high homology). Detailed post-annotation filtering strategy was described in detail previously^63^. For 14 tumor samples with paired normal WGS, germline SVs were removed followed by a second pass filtering against a normal in-house population database. For 8 tumor samples without paired normal WGS, SV filtration was performed using the population database removal method. Breakpoint microhomology and unbalanced/balanced translocation status was determined by *BRASS*.

SNV signature analysis was performed using *SigProfiler (v3.1)* R-package^68^. *PCAWG-HNSCC*, *PCAWG-BRCA1^mut^*, and *PCAWG-BRCA2^mut^* WGS cohorts^42^ were analyzed using an identical methodology to the Fanconi tumor WGS cohort.

Mouse tumor and pre-engraftment WGS samples were processed through the same workflow as human samples. Samples were aligned against *mm10* and somatic SV calls were made by filtration against pre-engraftment controls.

### Human Tumor HPV Detection

Reads from Fanconi SCC WGS and WES samples and PCAWG-HNSCC WGS samples not mapping to the human genome were aligned against 218 known HPV strain genomes (*NIH-PAVE* database)^69^ with *BWA-mem*. A sample was considered positive when a total of *≥* 1 unique paired-end reads aligned without clipping to a non-repetitive region of any HPV genome.

### Human Tumor Copy Number Alteration and Focal Peak Calling

Fanconi WES and WGS tumor samples were separately processed through the *CNVkit* (*v0.9.7*)^70^ pipeline to generate segmented copy-number alteration (CNA) profiles. Initial tumor CNA-ratio calls were made against a pooled-normal reference panel, comprised of either normal-WES samples (n=43) or normal-WGS samples (n=14) from our sequencing cohort. Circular binary segmentation was subsequently used to generate segmented CNA calls. Raw segmented CNA profiles from WGS and WES tumor cohorts were then combined for batch input into *GISTIC2* (v2.0.23)^71^. *GISTIC2* called recurrent focal amplification and deletion peaks in the Fanconi SCC cohort, including defining those genes contained within the boundaries of focal peaks. Variable peak calling parameters were set identical to the TCGA-HNSCC *GISTIC2* run (GDAC Firehose HNSCC run v2016_01_28).^72^ To account for variations in tumor purity affecting called CNA-depth, precomputed TCGA*-*HNSCC tumor purities (*Genomic Data Commons - TCGA Absolute Purity Master Calls)*^73^ and computed Fanconi SCC tumor purities (*Theta2 v0.7)*^74^ were used as inputs to normalize CNA signal amplitudes. *CNVkit* call tumor purity compensation function was employed for this transformation in both cohorts. Using the normalized CNA calls, high-level amplification (log2[CNA]*≥*0.9) and deletion (log2[CNA]*≤*0.9) thresholds were set for reporting significantly CN-altered genes within the defined focal peaks.

### Replication timing

DNA replication timing data was obtained from Koren et al., 2012^31^. The replication timing at each breakpoint of each CNA were calculated by linear interpolation of the replication timing values on the respective chromosomes. The distribution of replication timing of all breakpoints were then compared to the whole genome. P-values were calculated using the two-sample Kolmogorov-Smirnov test.

### Overlap test

A permutation-based overlap test was performed to assess whether FA SCC and PCAWG breakpoint sites were enriched for early replicating fragile sites^75^, common and rare fragile sites^76, 77^, and the aphidicolin breakome^78^. Breakpoints were stratified by type (deletion, inversion, translocation, or tandem duplication). Analysis was performed on each set of stratified regions compared to each set of fragile sites. Fragile site lists that were not originally in the hg19 genome were converted to hg19 using the UCSC LiftOver tool. In the case of early replicating fragile sites, gene locations were used for analysis. The fragile site lists were each permuted 1000 times by generating random genomic windows in the hg19 genome, with sizes equal to the original regions. These permuted windows were not permitted to overlap gaps in the reference genome, overlap each other, or overlap regions in the original gains/losses list. The occurrence of Fanconi anemia and PCAWG breakpoints in both permuted regions, and true fragile sites was determined. Then, a z-score was calculated for the number of breakpoints in true fragile sites compared to permutations, and a two-tailed p-value for observing a z-score as or more extreme was calculated.

### Human Tumor PacBio Sequencing Analysis

PacBio movies were converted to FASTQ (PacBio *bam2fastq*) and aligned to *hg19* using *NGMLR* (v0.2.7)^79^. SVs were called with *Sniffles* (v1.0.12)^79^. *Sniffles* detects deletions, tandem duplications, inverted duplications (BFBCs), inversions, translocations, and large insertions. The number of zero-mode waveguides (ZMWs) required to call an SV was set to 3 for 8 of 9 samples, ensuring that at least three native DNA molecules sharing the same breakpoint were present. One sample had below-average coverage, which required a ZMW threshold of 2 to prevent over-rejection. Raw calls were filtered using *AnnotSV*^80^ to remove population database germline SVs, including *GNOMAD*, *1000 Genomes Database*^81^, and *Database of Genomic Variants (DGV)*^82^, together representing approximately 15K human WGS samples and 10K CNV microarrays. We further removed SVs aligning proximal to centromeric and telomeric regions, as these were prone to spurious multiple-alignment calls. To account for germline SVs present in genomic regions inaccessible to short-read sequencing or arrays, an intra-cohort filtering step was applied to remove SVs co-occurring in more than one sample in our Pacbio cohort. SV breakpoint co-localization to known hg19 repeat regions was performed with *AnnotSV*, which subclassified repeats by *RepeatMasker*^83^ family grouping.

### Human Tumor 10X Linked-Read WGS Sequencing Analysis

Four Fanconi tumors were processed through the 10X *LongRanger* pipeline^84^. Barcoded clouds of linked-reads were aligned using *Lariat*^85^, followed by SNV/indel calling and haplotype-phasing with *Freebayes*.^86^ SVs were called using barcode-linkage, read-linkage, split-read evidence, and paired-end read support. Called somatic SVs have a minimum size threshold of 30kbp. SVs were filtered by polymorphic genomic region blacklisting in addition to calling confidence score. *Longranger* was run with the somatic flag enabled to increase the sensitivity of sub-haplotype event detection. Clusters of breakpoint-adjacent SVs sharing barcode overlap were outputted by the pipeline. These clusters were subsequently catalogued, validated, and manually reconstructed to define consensus linear SV chains.

### Algorithmic Detection and Chaining of Fold-Back Inversions, Templated Insertion Chains, and Retrotransposon Element Insertions from Illumina WGS

Each structural variant (SV) is characterized by two independent breakpoints and strand specificities by BRASS. Independent SVs are tested for being part of a composite genetic lesion whenever their breakpoints are proximal (distance *≤*2kb). Canonical inversions are recognized by two SVs with near-matching breakpoint positions and opposing strand specificities. Retrotransposon element (RTE) insertions are recognized by two independent SVs, most often translocations, where the majority of soft-clipped reads spanning the breakpoint in either region is characterized by long poly-A/poly-T tracts in the soft-clipped sequence. Only soft-clipped sequences of a minimal length of 20 bases were considered. Fold-back inversions (FBIs) were recognized as a single short-distance SV (*≤*2kb) with copy number alterations (CNA) emanating from the two breakpoint loci in a stepwise manner. FBIs with small templated insertions in a chain (TIC) were recognized as two independent proximal translocations, with local CNAs starting at the breakpoint loci, which upon tracing reveals a chain of small templated insertions (<= 1kb). Tracing was achieved by considering the supplementary alignment positions of the longest soft-clipped reads as the next step in the chain. There are two classes of FBI-TICs: (I) Cyclical FBI-TICs: tracing the chain from one breakpoint of the first proximal SV leads to the breakpoint of the other proximal SV after a few jumps in the chain (complete cycle) and (II) Broken FBI-TICs: tracing the chain from one breakpoint of the first proximal SV, or the second proximal SV, leads after a few jumps of the chain to the decoy contig, a region of extreme genomic homology (non-unique alignment) or the inability to recover the next step in the chain – the number of possible steps in the broken chain starting from the first and second proximal SVs are reported.

### Mouse and Human Tumor RNAseq Analysis

Transcript expressions were calculated using Salmon (v0.8.2)^87^ against mm10 and hg19. Gene expression levels as TPMs retrieved using Tximport (v1.8.0)^88^. Normalization and transformation of counts was performed using DESeq2 (v1.20.0).^89^ For comparison against TCGA-HNSCC RNAseq, raw FASTQ data was imported and run through the same pipeline. GSEA enrichment was performed using the GSVA (v1.34.0)^90^ and Limma (v3.42.2)^91^. Heat map generation was performed using Pheatmap R package (v1.0.10)^92^.

### Human Methylation Array Analysis

*Illumina EPIC Methylation* array beta-methylation signal scores were normalized and compared to TCGA-HNSCC *EPIC Methylation* beta-methylation array signal scores. We identified enriched/depleted genomic regions of methylation that were proximal or overlapping with known cancer-associated genes.

### Additional Software Used

Graphpad *PRISM (v*8) was used to generate scatter, bar, and pie charts, along with survival curves. *Maftools (v3.1.2)*^93^ R-package was used to generate the Fanconi SCC integrative oncoplot, Fanconi SCC/TCGA-HNSCC mutation comparison graph, and comparative SNV/Indel TMB plot. *Circa (v1.2.2)*^94^ was used to produce genomic circos plots. *SplitThreader (v1.0)*^95^ was used for supplementary visualization of PacBio SV events. *ASCAT*^96^ was used for generation of allele-specific copy number plots for select BMT-negative samples. *IGV (v2.8)*^97^ was used for manual assessment of structural variants.

## Code availability

https://github.com/MathijsSanders/SangerLCMFiltering

https://github.com/MathijsSanders/AnnotateBRASS

https://github.com/MathijsSanders/ExtractSVChains

---

**Extended data Fig.1.**
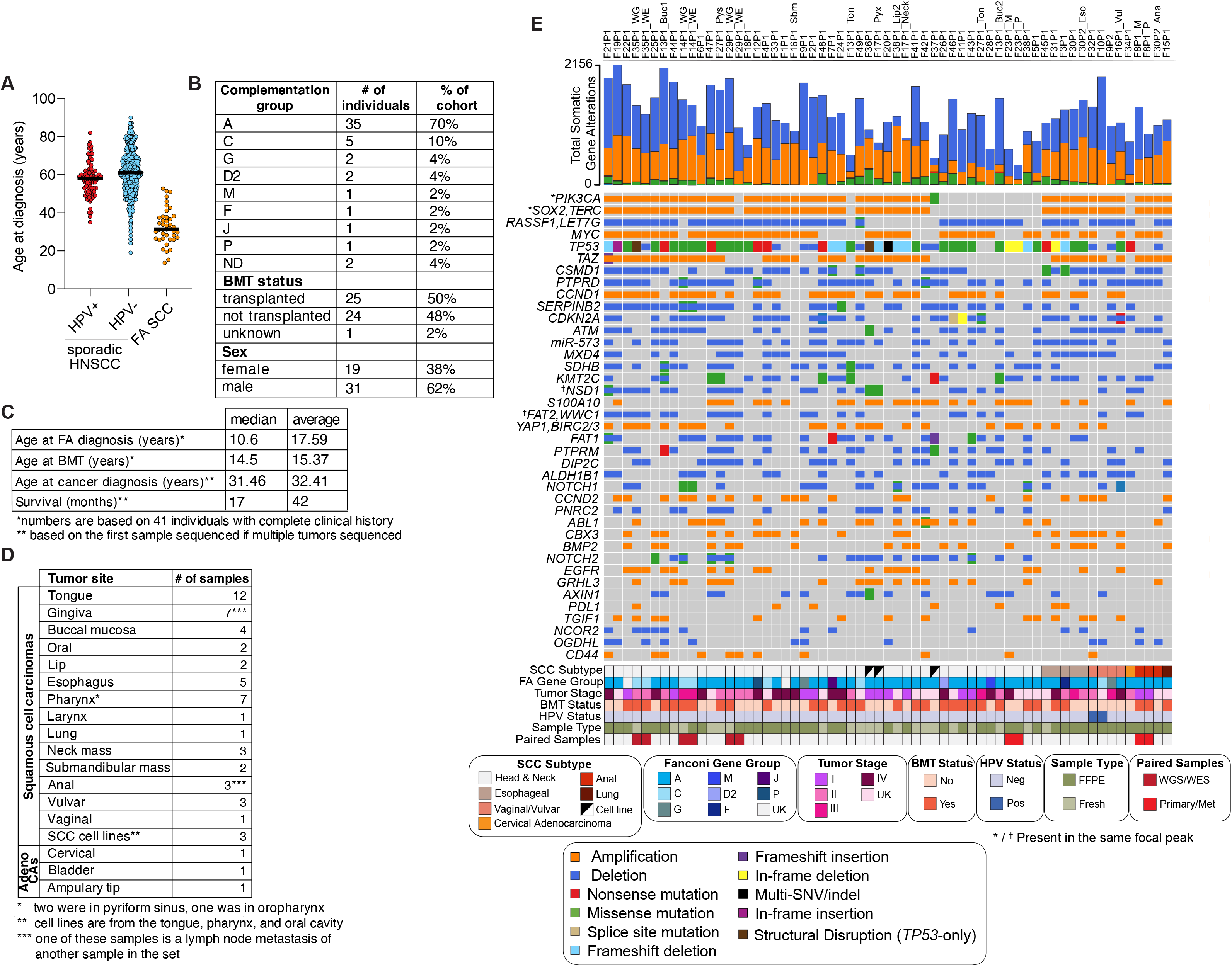
Clinical characteristics of the FA SCC cohort and mutational landscape of individual FA SCCs. **a** Age at diagnosis of individuals in the FA SCC (n=41), HPV+ (n=78) and HPV- (n=419) sporadic HNSCC cohorts, for which full clinical data was available. Clinical data for sporadic HNSCC cohorts were obtained from TCGA database. **b** and **c** Characteristics of 41 FA individuals with complete clinical information. Some individuals had multiple cancers. For these cases, survival was calculated from the first cancer sequenced in this study. **d** Type and site of the sequenced tumors. **e** Oncoplot of the FA SCC cohort indicating type of variant affecting the genes indicated on the left. Recurrent focal CNAs were defined by *GISTIC2*. Amplifications were classified as log2(sCNA)≥0.9 and focal deletions as log2(sCNA)≤-0.9 after normalizing for tumor purity. Samples are stratified by SCC tissue subtype. One adenocarcinoma sample (cervical adenocarcinoma) is shown, while the bladder and intestinal adenocarcinomas are not displayed.

**Extended data Fig. 2.**
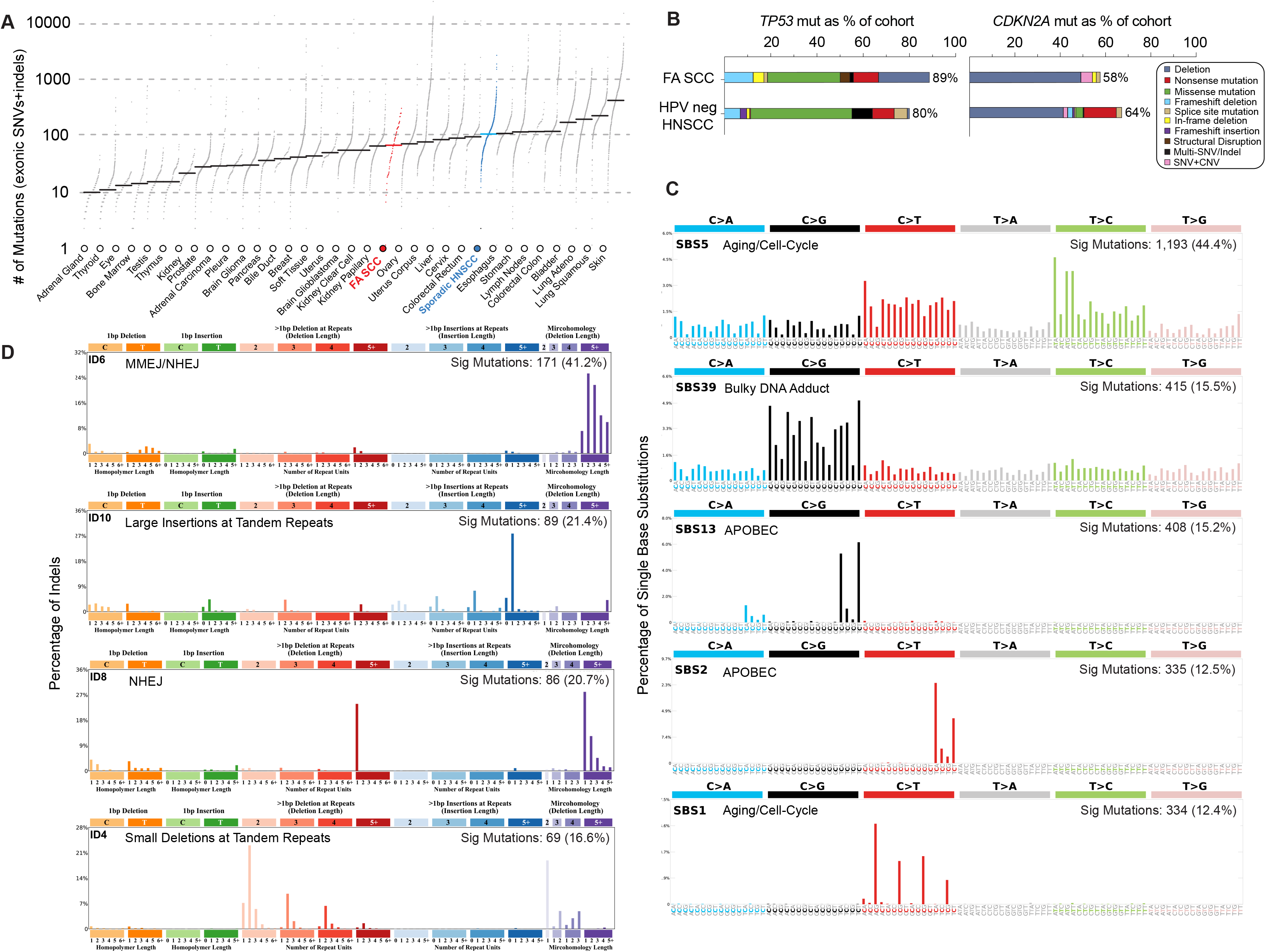
Characterization of the SNV burden and COSMIC SNV mutation signatures in FA SCCs. **a** Comparison of tumor mutation burden (TMB) between TCGA cohorts and FA SCC cohort (n=55 independent SCCs). Each dot represents the number of exonic SNV and indel mutations per sample, with median mutation rate indicated with a black horizonal line. FA SCC samples are colored red and TCGA-HNSCC samples are colored blue. **b** Mutation type frequency in *TP53* and *CDKN2A* genes in the FA SCC and HPV-negative TCGA-HNSCC cohorts. **c** *SigProfiler-*extracted de-novo COSMIC SNV (SBS) signatures in FA SCC cohort, calculated from 16 independent samples with >100 somatic SNV events and low to no detectable BMT donor SNP contamination. **d** *SigProfiler-*extracted de-novo COSMIC indel (ID) signatures in FA SCC cohort, calculated from 55 independent FA SCC samples.

**Extended data Fig. 3.**
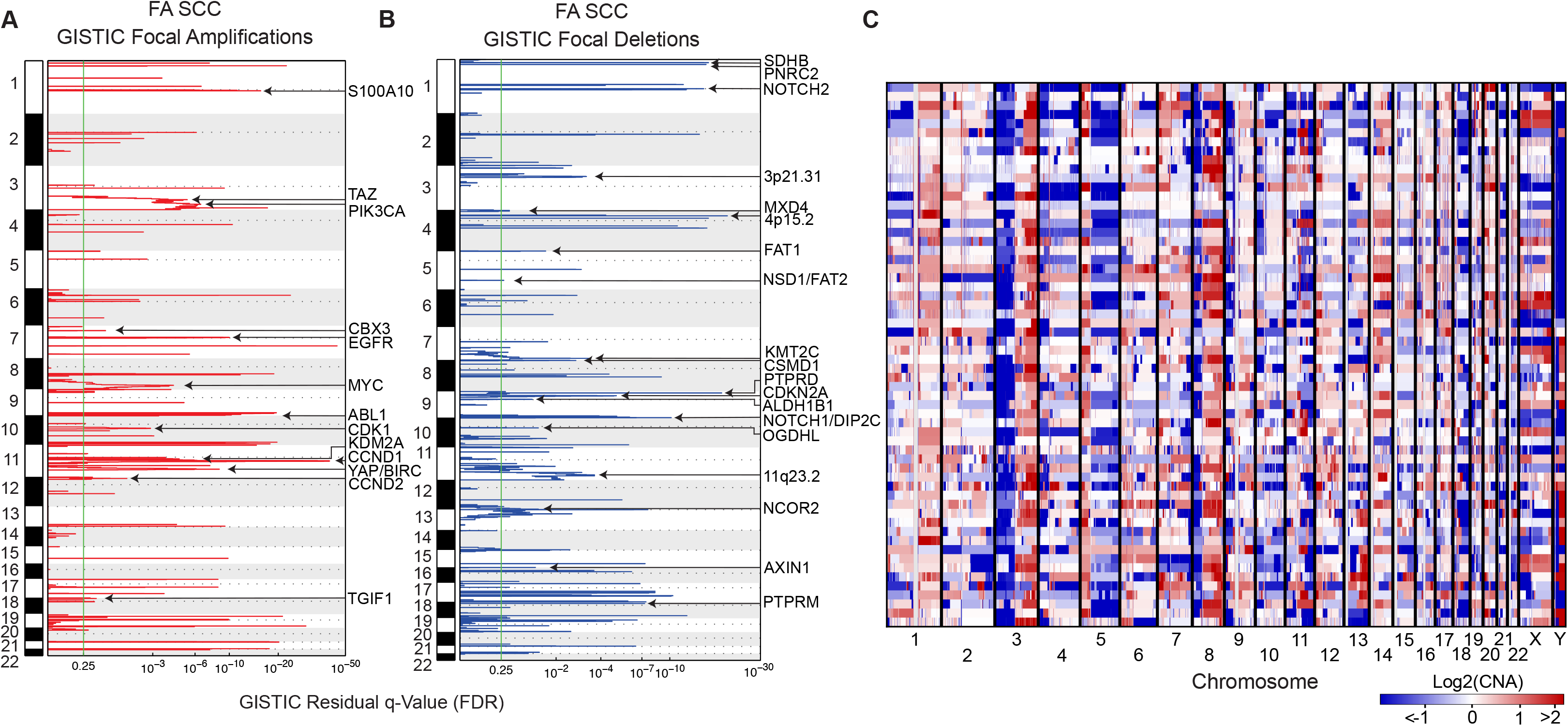
Copy number instability in FA SCCs. **a** Plot displaying chromosomal locations of recurrent focal amplification peaks detected by *GISTIC2* in all FA SCC samples (n=60 samples, including 55 independent SCCs, 2 SCC metastases, and 3 SCC samples sequenced by both WGS/WES) and one cervical adenocarcinoma. GISTIC residual q-value is shown below, with default minimum calling threshold displayed as green line. **b** A plot displaying chromosome location of recurrent focal deletion peaks detected by *GISTIC2* in all Fanconi SCC samples (n=60). GISTIC residual q-value is shown below, with default minimum calling threshold displayed as green line. **c** Genomic copy-number alteration heatmap displaying individual tumor sCNAs in all FA SCC samples (n=60) and one cervical adenocarcinoma, coloured by amplitude intensity and normalized for individual tumor purity. Each row is a tumor sample.

**Extended data Fig. 4.**
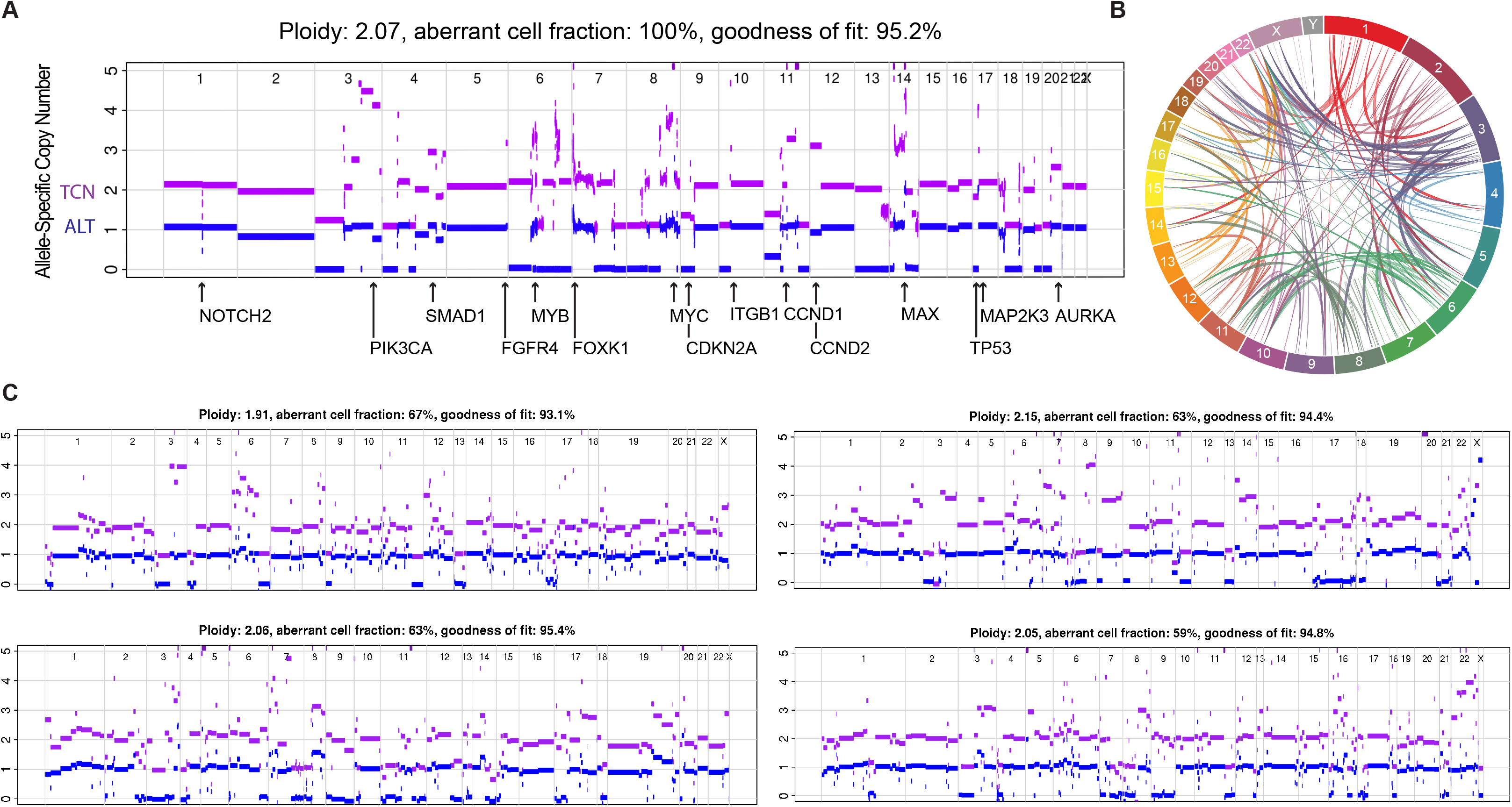
Allele-Specific Copy Number Analysis of Tumors (ASCAT) profiles of FA SCCs from individuals who have not undergone a hematopoietic stem cell transplantation. **a** *ASCAT* plot of a WGS-sequenced FA SCC sample (F17P1). Total copy number is represented by a purple line. Minor allele is represented by a blue line. Indicated are notable oncogenes and tumor suppressors localizing to focal CNA regions. **b** Genomic circos plot displaying all somatic SV events detected from Illumina WGS in F17P1 sample depicted in panel a. **c** *ASCAT* plots displaying allele-specific copy number alteration in select WES-sequenced FA SCC tumors with little to no detectable BMT-donor SNP contamination. Upper left (F32P1), Upper right – (F16P1-Vulv), Bottom left (F4P1), Bottom right (F25P1). F32P1 is HPV+, but harbours somatic deletions of *TP53* and *CDKN2A*.

**Extended data Fig. 5.**
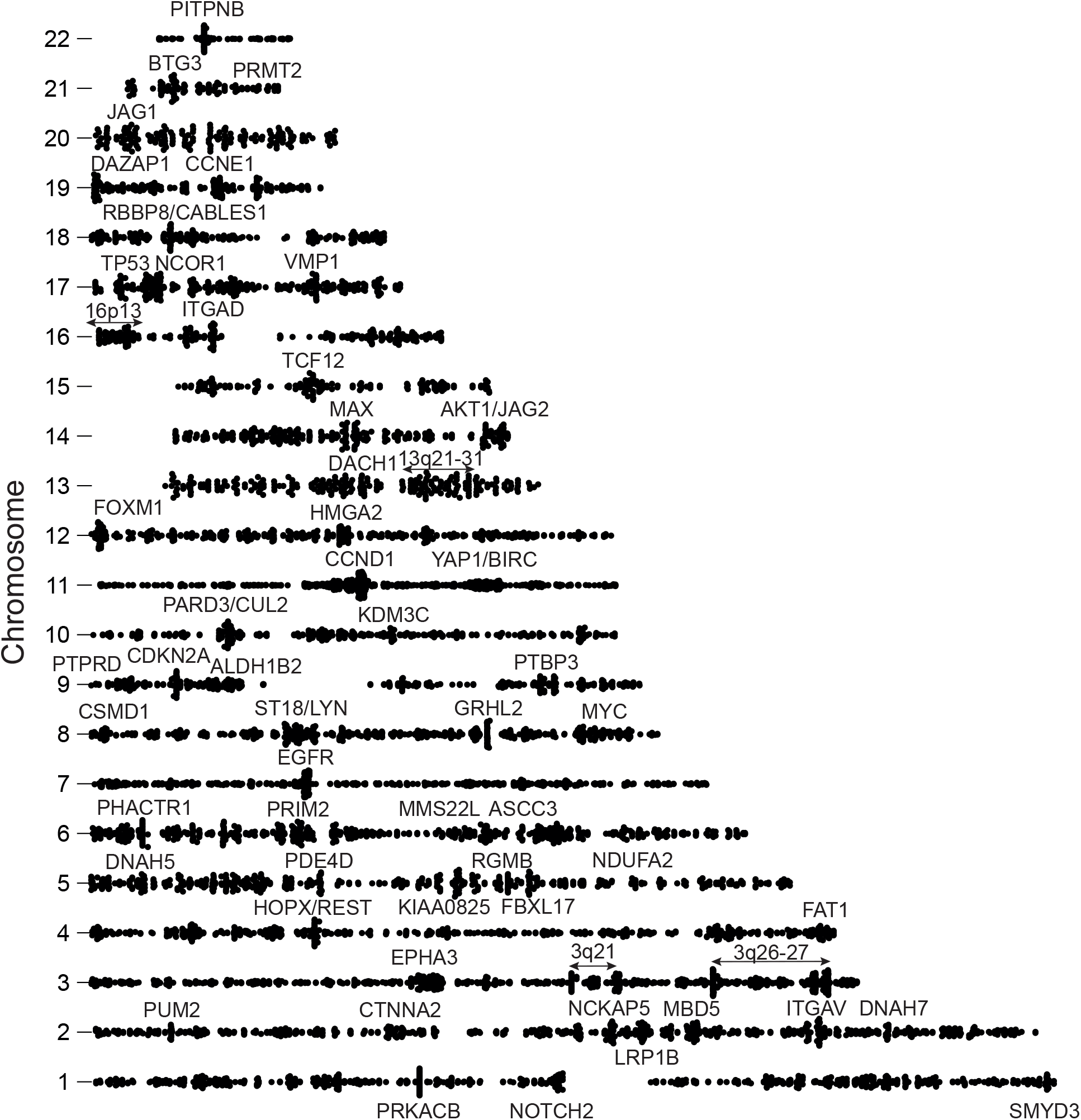
Characterization of SVs and their breakpoints in FA SCCs. **a** Scatter plot displaying localization of 8,896 SV breakpoints (from 4,448 SVs) in FA SCC by chromosome and genomic position. Relative breakpoint density is indicated by height off the baseline. Annotated are curated oncogenes and tumor suppressors localizing to breakpoint hotspots.

**Extended data Fig. 6.**
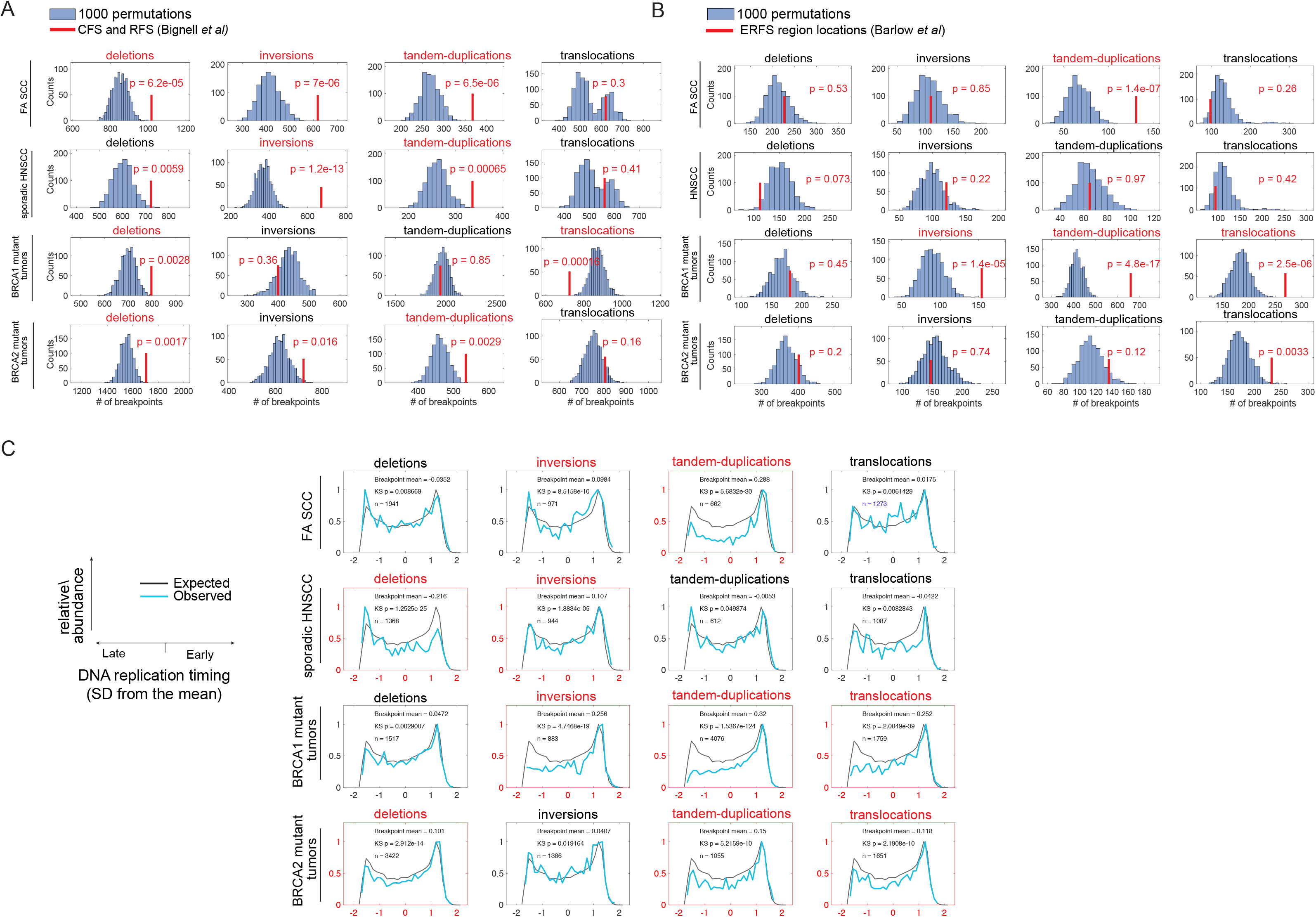
Localization of SV breakpoints from FA SCC, sporadic HNSCC, BRCA2^mut^, and BRCA1^mut^ tumors relative to genome replication timing and fragile sites. **a** Binned count of the SV breakpoints from the indicated cohorts and SV class, localizing to common and rare fragile sites. SV class highlighted in red indicates a significant association, as determined by two-tailed p-value of z-score compared to 1000 permutations of fragile site locations. **b** Binned counts of the SV breakpoints localizing to “early-replicating fragile sites”. SV class highlighted in red indicates a significant association, as determined by two-tailed p-value of z-score compared to 1000 permutations of fragile site locations **c** Replication timing of the SV breakpoint loci. Plotted in black is the expected replication timing distribution. Plotted in blue is the observed localization of SV breakpoints to early, mid, or late replicating genomic regions. Vertical axis indicates relative abundance and horizontal axis indicates standard deviation from mean replication timing. Kolmogorov-Smirnov (KS) significance values are indicated. *n* corresponds to the number of breakpoints included in the sample for each analysis.

**Extended data Fig. 7.**
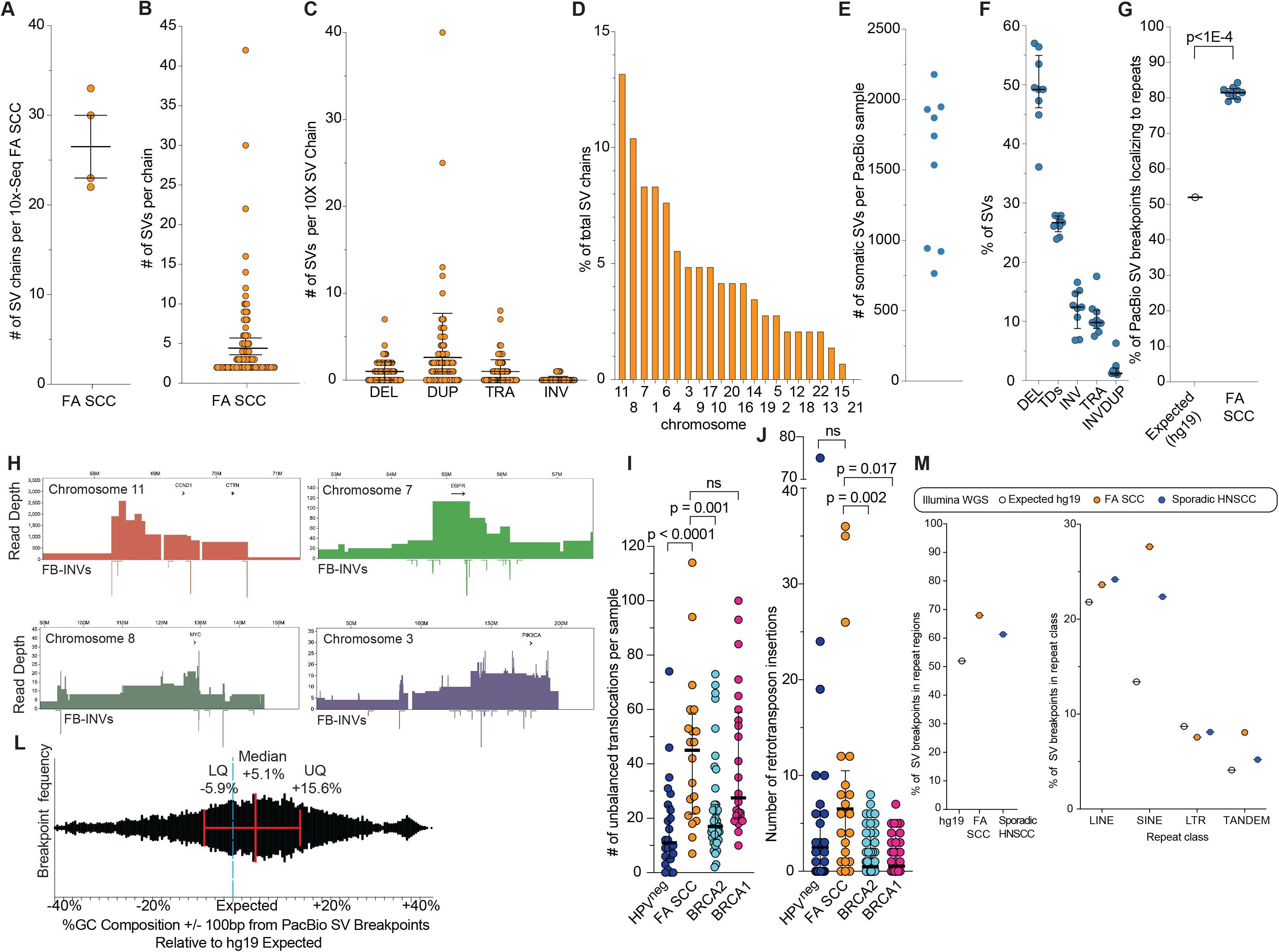
Further analysis of complex structural variants in FA SCC assessed using 10x linked read sequencing, PacBio long read sequencing, and lllumina WGS. **a** Number of somatic SV chains detected in 10X-sequenced Fanconi SCC samples (n=4), where a chain is defined as ≥ 2 discrete SVs (≥ 4 unique breakpoints). Median and IQR is indicated. **b** Number of SVs present in 108 SV chains in 10X-sequenced FA SCCs. Mean (4.6 SVs) and IQR is indicated. **c** Number of SVs of indicated class present in 108 SV chains from 10X-sequenced FA SCCs. Means and IQRs are indicated. **d** Distribution of SV breakpoints from 108 SV chains according to human chromosome number. **e** Number of somatic SVs identified in 9 PacBio-sequenced FA SCCs. 3 samples were sequenced to 10x average coverage, and 6 samples were sequenced to 30x average coverage. **f** Proportion of somatic SVs represented by each SV class in 9 PacBio-sequenced FA SCCs. Medians and IQRs are indicated. **g** Comparison of hg19 expected vs observed percentage of somatic SV breakpoints in 9 PacBio-sequenced FA SCCs that localize to global repeat regions. Mann-Whitney U test two-tailed exact p-values is indicated, with median and IQR shown. **h** Examples of fold-back inversions (FBI) driving sharp copy-number change at key oncogenic loci identified in FA SCCs (PacBio data). **i** Comparison of the raw number of unbalanced translocation events in FA SCC (n=20), HPV-negative sporadic HNSCC (n=24), BRCA2^mut^ (n=41), and BRCA1^mut^ (n=24) cohorts. Mann-Whitney U test two-tailed exact p-values are indicated, with median and IQR shown. **j** Comparison of the number of retrotransposon element (RTE) insertions in FA SCC (n=20), HPV-negative sporadic HNSCC (n=24), BRCA2^mut^ (n=41), and BRCA1^mut^ (n=24) cohorts. Mann-Whitney U test two-tailed exact p-values are indicated, with median and IQR shown. **k.** Breakpoint density graph displaying GC% sequence composition within +/− 100bp from SV breakpoints identified in PacBio sequencing, calculated relative to hg19 global GC% frequency (40.9%) (notated as “expected”). Median, upper, and lower quartiles are displayed. **l.** Comparison of hg19 expected vs. observed percentage of somatic SV breakpoints from FA SCCs (n=20) and sporadic HNSCCs (n=24) that localize to repeat regions of the human genome and to the indicated repeat class (Illumina WGS).

**Extended Fig. 8.**
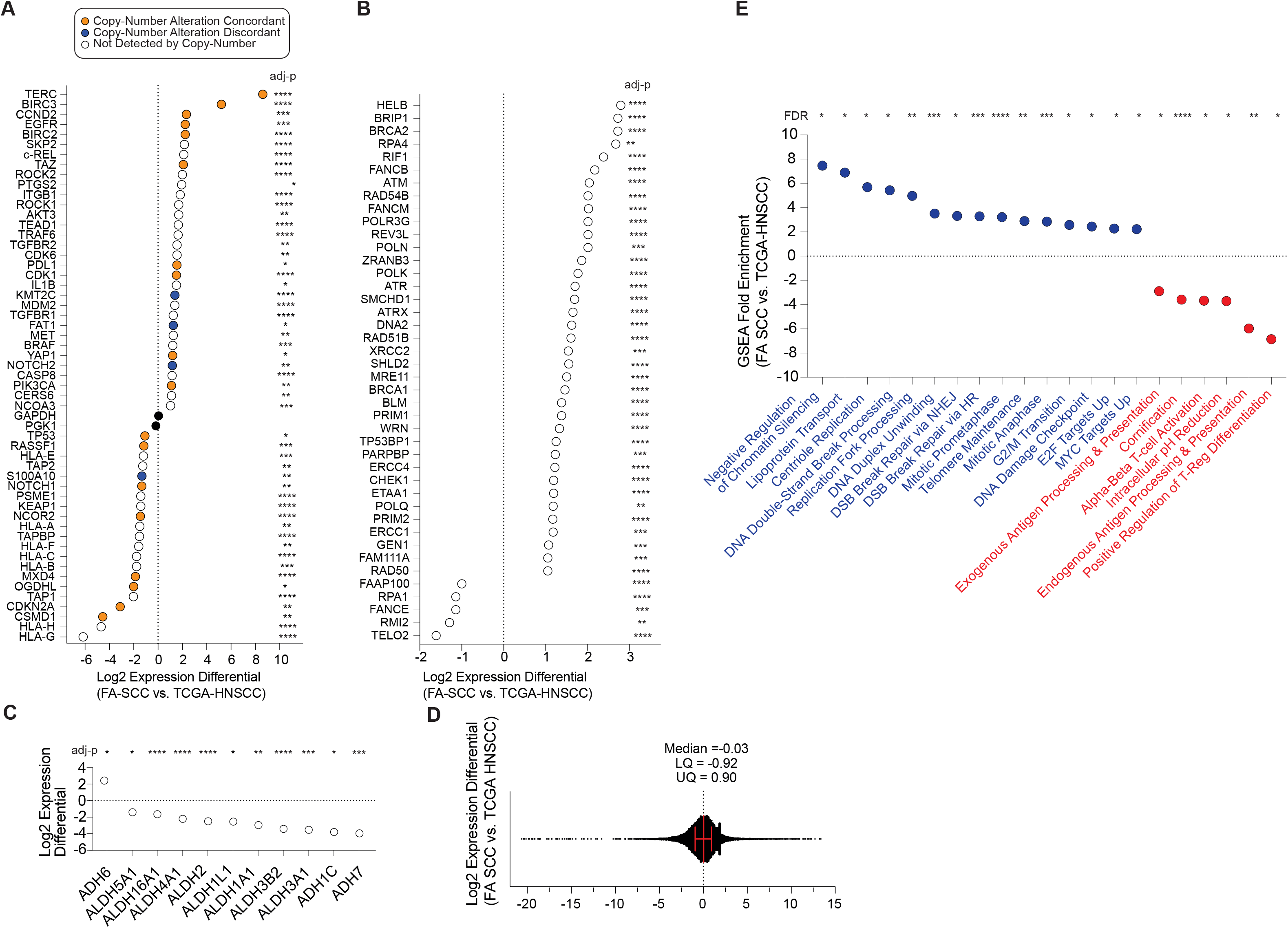
Comparison of gene expression between FA SCC and sporadic HNSCC. **a** Cancer-relevant genes differentially expressed between FA SCC (n=6 tumors) and sporadic HNSCC (n=520 tumors) as assessed by RNAseq, including genes displayed in Fig. 1C. Differential expression is gated at log2(FC)>1 or log2(FC)<-1 with DESeq2 adjusted p-value <0.05. Genes whose relative expression is supported by sCNA are colored orange. Genes whose relative expression is discordant with sCNA frequency are colored blue. Genes not identified in focal CNA peaks are colored white. *GAPDH* and *PGK1* are indicated in black as housekeeping controls. **b** DNA repair genes differentially expressed in FA SCC (n=6) versus sporadic HNSCC (n=520) by RNAseq. Differential expression is gated at log2(FC)>1 or log2(FC)<-1 with DESeq2 adjusted p-value <0.05. **c** Aldehyde dehydrogenase (Aldh) and alcohol dehydrogenase (Adh) genes differentially expressed between FA SCC (n=6) and sporadic HNSCC (n=520). Differential expression is gated at log2(FC)>1 or log2(FC)<-1 with DESeq2 adjusted p-value <0.05 **d** Quality-control distribution graph showing log2(FC) values of all genome-wide transcripts comparing FA SCC (n=6) vs sporadic HNSCC (n=520). **e** Gene-set enrichment/depletion (GO) analysis of genes differentially expressed between FA SCC and sporadic HNSCC. Genes entered into analysis were gated at log2(FC)>1 or log2(FC)<-1 with DESeq2 adjusted p value <1E-5. Gene sets were gated at >2-fold enrichment over expected background with adjusted p-value <0.01 to be reported in figure.

**Extended Fig. 9.**
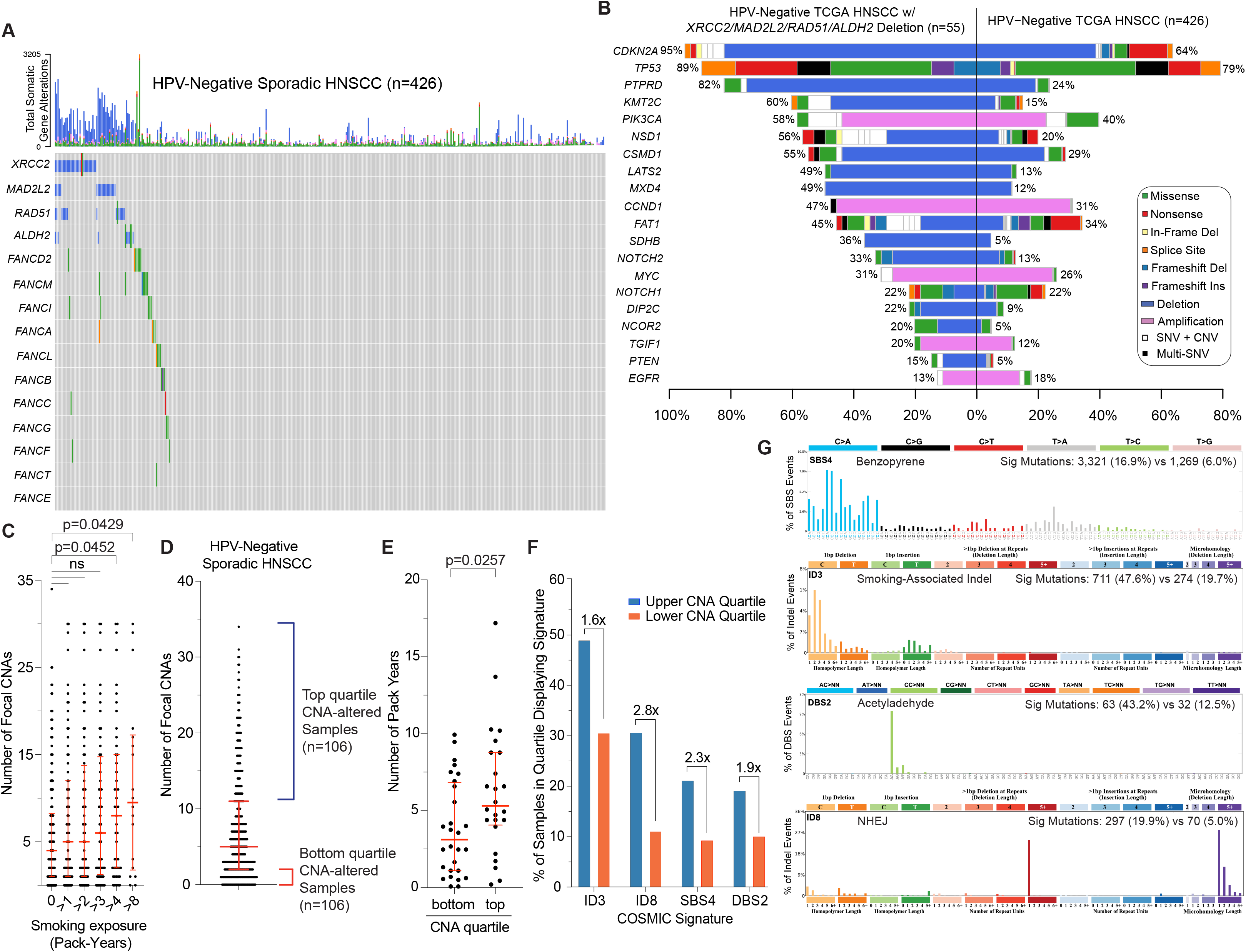
Copy-number instability in sporadic HNSCC and FA pathway deficiency. **a** Oncoplot of 426 HPV-negative sporadic HNSCC tumors with somatic copy-number alteration (sCNA) or SNV/indel-alteration of Fanconi pathway genes or *ALDH2.* Mutation type is indicated in the legend. Top bar graph indicates the relative copy-number instability of each sample. Blue indicates deletions, magenta indicates amplifications. **b.** Mutational frequency of key HNSCC driver genes in HPV-negative sporadic HNSCC samples with *MAD2L2*, *ALDH2*, *RAD51*, or *XRCC2* deletions (n=55) versus entire HPV-negative TCGA-HNSCC cohort (n=426). **c** Number of focal copy-number alterations in HPV-negative sporadic HNSCC tumors (n=426), stratified by number of clinically annotated cigarette pack-years associated with each patient sample. Shown are cases with zero pack years (no recorded smoking), cases with more than one (>1) pack-years, and cases with more than two (>2), more than three (>3), more than four (>4) and more than 8 (>8) pack years. Mann-Whitney U test two-tailed exact p-values are indicated, with median and IQR shown. **d** HPV-negative sporadic HNSCC samples (n=426) ranked by number of focal somatic copy-number alteration (sCNA) peaks as defined by *GISTIC2*. Annotated are the top and bottom sCNA quartiles, with the top being most unstable and the bottom being most stable. **e** Comparison of the number of cigarette pack-years for smokers in top and bottom copy-number quartiles. Mann-Whitney U test two-tailed exact p-value is indicated, with median and IQR shown. **f** Bar chart indicating the proportion (%) of samples within top and bottom sCNA quartiles exhibiting at least 3 mutational occurrences of each respective COSMIC signature ID3, ID8, SBS4, or DBS2. Annotated are fold-differences in these proportions. **g** Comparison of the total number of ID3, ID8, SBS4, and DBS2 signature events between top and bottom sCNA quartiles. Indicated in brackets is the proportion (%) of total SBS, DBS, or ID events represented by the respective signature in each sCNA quartile.

**Extended Fig 10.**
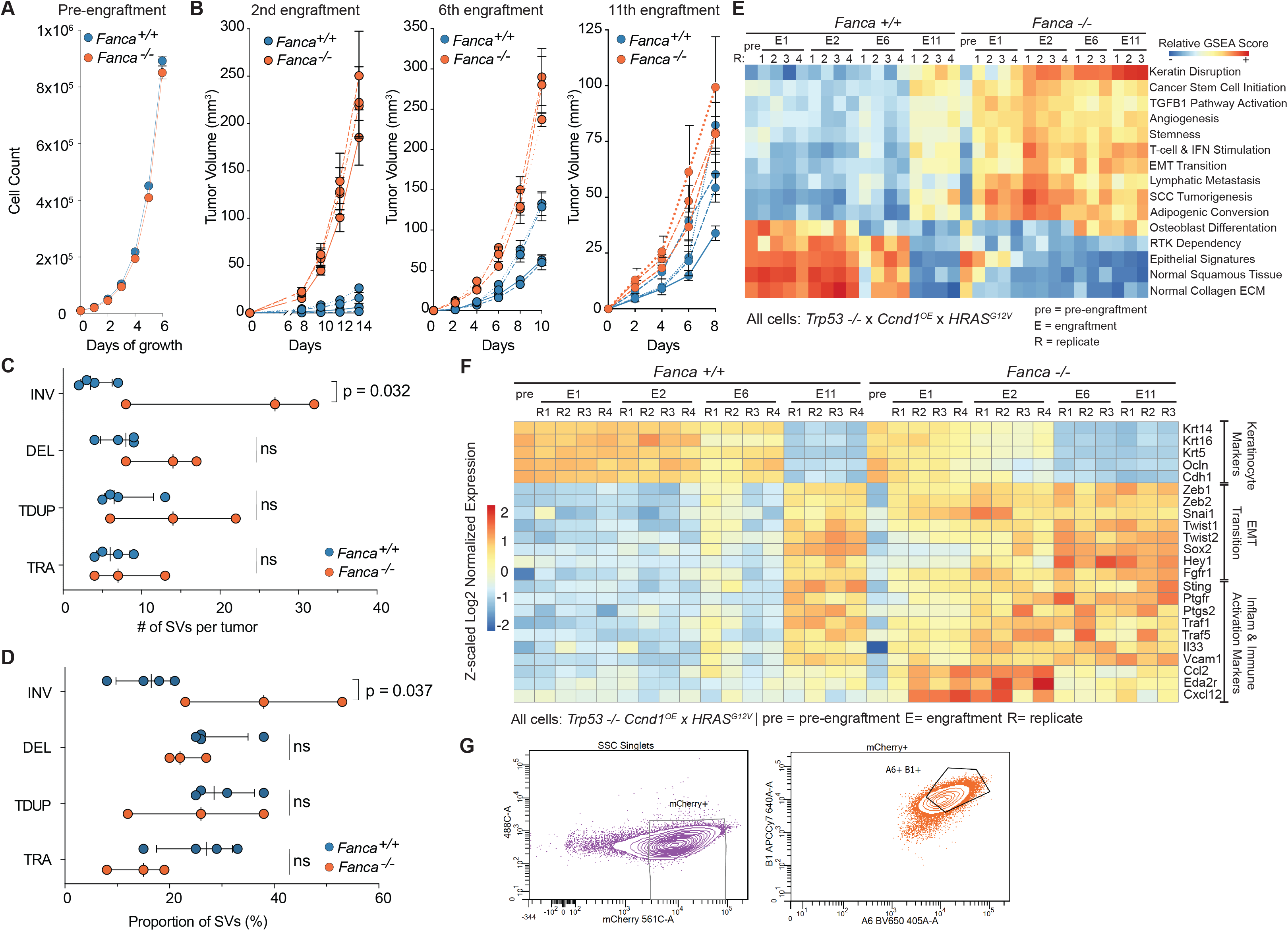
Characterization of a murine FA SCC model. **a** *In vitro* cell growth curve of pre-engraftment *Fanca^+/+^* and *Fanca^−/−^* keratinocytes, measured by cell count over six days with five replicates per genotype. Data points indicate the mean cell count and bars indicate standard error. **b** Mean replicate tumor volumes measured at multiple time points during the 2^nd^, 6^th^, and 11^th^ engraftment cycles of *Fanca^+/+^* and *Fanca^−/−^* keratinocytes. Each genotype has 4 independent replicates, each of which in turn is comprised of 4 co-engrafted tumor sites on a single mouse (for a total of 16 tumors per genotype). Each data point represents one replicate as the mean volume of its 4 constituent tumors at the specified time point, with standard error bars indicated. 100×10^3^, 70×10^3^, and 35×10^3^ cells were engrafted at 2^nd^, 6^th^, and 11^th^ engraftment respectively. 1^st^ engraftment data is shown in Fig. 4C. Fanca^−/−^ was reduced to 3 replicates at the 6^th^ and 11^th^ cycles due to replicate loss from host death. **c** Number of tumor SVs categorized by class: inversion (INV), deletion (DEL), tandem duplications (TD), translocation (TRA) in 4 *Fanca^+/+^* and 3 *Fanca^−/−^* replicates from 6^th^ engraftment cycle. Two-tailed, unpaired t-test p values displayed, with medians and IQRs indicated. **d** Proportion (%) of SVs represented by each class in *Fanca^+/+^* and *Fanca^−/−^replicates* at 6^th^ engraftment cycle. Two-tailed, unpaired t-test p values displayed, with medians and IQRs indicated. **e** Unsupervised-clustering heatmap displaying differential transcriptomic gene-set enrichment across all replicates at pre-engraftment and 1^st^, 2^nd^, 6^th^, & 11^th^ engraftment cycles for *Fanca^+/+^* and *Fanca^−/−^* genotypes. Relative gene set enrichment or depletion is indicated by color scale at each time point (ANOVA test). Gene sets displayed have an adjusted p value < 0.0000001. Pre indicates pre-engraftment, E indicates engraftment, R indicates replicate. **f** RNAseq differential expression heatmap across all replicates displaying time-course expression changes in genes associated with keratinocyte identity, EMT transition, and inflammation/immune cell activation. Heatmap color indicates Z-scaled log2-normalized expression. Pre indicates pre-engraftment, E indicates engraftment, R indicates replicate. **g** Flow-cytometry gates for integrin a6 (BV650), integrin b1 (APC/Cy7), and Ccnd1 (mCherry) during mouse keratinocyte line generation.

